# Disease-associated microglia and activation of CD8^+^ T cells precede neuronal cell loss in a model of hereditary spastic paraplegia

**DOI:** 10.1101/2024.09.02.610538

**Authors:** A Frolov, H Huang, D Schütz, M Köhne, N Blank-Stein, C Osei-Sarpong, M Büttner, T Elmzzahi, M Khundadze, M Becker, L Bonaguro, CA Hübner, K Händler, R Stumm, E Mass, M Beyer

**Affiliations:** Immunogenomics & Neurodegeneration, Deutsches Zentrum für Neurodegenerative Erkrankungen (DZNE) e.V., 53127 Bonn, Germany; Systems Medicine, Deutsches Zentrum für Neurodegenerative Erkrankungen (DZNE) e.V., 53127 Bonn, Germany; Department of Microbiology and Immunology, The Peter Doherty Institute for Infection and Immunity, University of Melbourne, Melbourne, VIC, Australia; Developmental Biology of the Immune System, Life & Medical Sciences (LIMES) Institute, University of Bonn; 53115 Bonn, Germany; Institute of Pharmacology and Toxicology, Jena University Hospital, Friedrich-Schiller-University Jena, 07747 Jena, Germany; Institute of Experimental Pathology, Centre of Molecular Biology of Inflammation, University of Münster, Von-Esmarch-Esmarch-Str. 56 48149 Münster, Germany; Genomics & Immunoregulation, Life & Medical Sciences Institute, University of Bonn, Bonn, Germany; Institute of Human Genetics, Jena University Hospital, Friedrich-Schiller-University Jena, 07747 Jena, Germany; Center for Rare Diseases, University Hospital Jena, Friedrich-Schiller-University, Am Klinikum 1, 07747 Jena, Germany; Modular High-Performance Computing and Artificial Intelligence, Deutsches Zentrum für Neurodegenerative Erkrankungen (DZNE) e.V., 53127 Bonn, Germany; Institute of Human Genetics, Universitätsklinikum Schleswig-Holstein, University of Lübeck and University of Kiel, 23562 Lübeck, Germany; PRECISE Platform for Single Cell Genomics and Epigenomics, DZNE and University of Bonn, 53127 Bonn, Germany

## Abstract

In central nervous system (CNS) diseases characterized by late-onset neurodegeneration, the interplay between innate and adaptive immune responses remains poorly understood. This knowledge gap is amplified by the prolonged nature of these diseases, complicating the delineation of brain-resident and infiltrating cells. Here, we conducted a comprehensive profiling of innate and adaptive immune cells across various CNS regions in a murine model of spastic paraplegia 15 (SPG15), a complicated form of hereditary spastic paraplegia (HSP). Using fate-mapping of bone marrow-derived cells via genetic labeling, we identified microgliosis and microglial MHC-II upregulation accompanied by infiltration and local expansion of T cells in the CNS of *Spg15^-/-^* mice. Single-cell analysis revealed an increase of disease-associated microglia (DAM) and clonal expansion of effector CD8^+^ T cells across CNS regions occurring prior to neuronal loss. Analysis of potential cell-cell communication pathways suggested bidirectional interactions between DAM and effector CD8^+^ T cells potentially contributing to disease progression in *Spg15^-/-^* mice. In summary, we identified a shift in microglial phenotypes associated with recruitment and clonal expansion of T cells as a new characteristic of *Spg15*-driven neuropathology. Targeting activated microglia, CD8^+^ T cells and their communication represent promising avenues to prevent the loss of neuronal function in HSP.

## Introduction

Hereditary spastic paraplegias (HSPs) constitute a heterogeneous group of genetic disorders characterized by corticospinal tract dysfunction, leading to progressive gait disturbance and lower limb spasticity (Fereshtehnejad et al., 2023; McDermott et al., 2000). Over 80 gene loci designated as spastic paraplegia genes (*SPGs*) have been identified as causative factors for HSPs, with genes implicated in various cellular functions such as membrane trafficking, mitochondrial function, nucleotide and lipid metabolism, organelle biogenesis as well as myelination (Blackstone, 2018).

A prominent neuropathological feature of HSPs is axonal degeneration affecting corticospinal and ascending tracts. Although loss of cortical layer V motor neurons may occur in HSPs, it is likely to be secondary to axonal damage (McDermott et al., 2000). Recent work employing murine HSP models underscore the role of neuroinflammation in HSP. Specifically, mice with mutations in the myelin-related proteolipid protein-1 gene (*Plp1*), which corresponds to *Spg2*, exhibit disease-amplifying neuroinflammation (Groh et al., 2016). This observation was extended to a murine model of HSP type 11 (*Spg11*) where deletion of *Spg11,* the gene encoding Spatacsin, which interacts with the fifth adaptor protein complex (AP5) involved in endosomal traffic, lysosomal biogenesis, and autophagy (Pozner et al., 2020), recapitulated the motor phenotype of SPG11 patients, revealing microgliosis and CD8^+^ T cell accumulation in the brain (Hörner et al., 2022; Branchu et al., 2017; Varga et al., 2015).

In this study, we investigated neuroinflammation in a murine model of HSP type 15 (*Spg15^-/-^*) (Khundadze et al., 2013) mimicking a disease-causing *SPG15* germline loss-of-function mutation causing disease in humans (Saffari et al., 2023). *SPG15* (*ZFYVE26*) encodes for Spastizin, an interaction partner of Spatacsin. Despite operating in the same cellular processes, Spatacsin and Spastizin exhibit distinct functions in autophagy and endocytosis (Vantaggiato et al., 2019; Khundadze et al., 2021). Similar to *Spg11*^-/-^ mice, *Spg15*^-/-^ mice manifest progressive motor deficits and loss of cortical layer V motoneurons and cerebellar Purkinje cells by 16 months of age (Khundadze et al., 2013). *Spg15*^-/-^ mice thus represent a model of slowly developing neuronal dysfunction with late-onset neurodegeneration that phenocopy the human disease pathology. SPG15 and SPG11 patients are clinically indistinguishable and represent the largest part of complicated HSP cases characterized by childhood mental impairment, followed by the onset of walking difficulties and progressive spasticity during the 2nd or 3rd decades (Pensato et al., 2014). Intriguingly, SPG15 patients often exhibit a background of speech delay and learning disability preceding the onset of the progressive motor symptoms by several years (Saffari et al., 2023). Yet, the mechanisms underlying the impaired motor and higher-order brain functions in complicated HSPs are still poorly understood.

Given the growing evidence of neuroinflammatory processes contributing to the pathobiology of various HSP types (Groh et al., 2016; Branchu et al., 2017; Hörner et al., 2022), an in-depth single-cell characterization of microglia and T cells is needed. This approach is of particular interest since a fine-tuned transcriptional differentiation and functional specialization has been described for microglia and CD8^+^ T cells in ageing and several neurodegenerative diseases (Kaya et al., 2022; Chen and Colonna, 2021; Chen et al., 2023). The advent of single-cell transcriptomics has allowed the identification of transcriptionally defined cell states, including the disease-associated microglia (DAM) program, which is associated with microglial activation during neuroinflammation and neurodegeneration (Deczkowska et al., 2018; Paolicelli et al., 2022; Keren-Shaul et al., 2017).

Microglial activation can lead to an influx of adaptive immune cells, perpetuating neuroinflammation. While earlier studies suggested that CD8^+^ T effector memory (Tem) cells are anti-inflammatory in the aging CNS by supporting homeostatic microglial functions (Ritzel et al., 2016), recent findings indicate that clonally expanded interferon-gamma (IFNγ)^+^/PD1^+^ CD8^+^ T cells negatively affect neural stem cells (Dulken et al., 2019). In line with a detrimental role, CD8^+^ T cells were implicated in the formation of IFN-responsive microglia in aging white matter (Kaya et al., 2022). Moreover, clonally expanded activated CD8^+^ T cells worsen tau pathology by driving microglia to a DAM-like state (Chen et al., 2023).

In late-onset neurodegenerative disorders, adaptive immune responses and interactions between innate and adaptive immune cells have been neglected. Furthermore, the extent of local expansion of CNS-resident immune cells early in life and immune cell recruitment during disease progression remain largely unclear. To address this, we established *Cxcr4^CreERT2^* in combination with *Rosa26^tdTomato^* as a tool to permanently label long-term hematopoietic stem cells (HSC) and unequivocally track HSC progeny in the inflamed brain (Werner et al., 2020). Using the combination of *Cxcr4^CreERT2^*-mediated fate mapping in *Spg15*^-/-^ mice with immunohistology and multi-omic analysis of microglia and T cells, we provide the first comprehensive assessment of the CNS immune response in a model of complicated HSP.

## Results and discussion

### Microglia activation precedes neuronal cell loss in *Spg15^-/-^* mice

A previous study of 15 months *Spg15^-/-^* mice indicated the presence of astrogliosis across different brain regions (Marrone et al., 2022). Since microglial activation precedes astrogliosis in the context of neuroinflammation (Liddelow et al., 2017), we hypothesized that *Spg15* deficiency may result in immune cell activation, thereby contributing to neurodegeneration. To address a possible contribution of immune cells to neuronal loss observed in 15 months *Spg15^-/-^* mice (Marrone et al., 2022), we first quantified neuronal cell numbers in the spinal cord (SC) and cortex of younger animals. At 12-15 months of age, we could not detect a difference between *Spg15^-/-^* mice and littermate controls in the abundance of NeuN^+^ cells in the ventral horn of the lumbar SC and layers V / VI of the primary motor cortex (Fig. 1A, B), despite an obvious gait phenotype in *Spg15^-/-^* mice at this age (Fig. 1C) indicative of a slowly progressing neuronal dysfunction (Khundadze et al., 2013; Marrone et al., 2022). Since many neurodegenerative diseases are accompanied by neuroinflammation and microglial activation (Paolicelli et al., 2022), we next assessed the activation of microglia by immunofluorescence stainings. Our analysis indicated an increase of Iba1 signal in SC and cortex, which was accompanied by an increase of Iba1^+^ cell numbers (Fig. 1D, E). In the SC, these changes were most prominent in white matter but were also noted in grey matter. In the brain, multiple regions were affected, including the motor cortex, sensory regions, such as the dorsal thalamus, and regions involved in cognition, like the hippocampus. These qualitative observations were substantiated by quantification of Iba1 and P2ry12 signals in the ventral SC and motor cortex (Fig. 1D-G). Given that Iba1 is increased and P2ry12 decreased in reactive microglia (Mildner et al., 2017; Haynes et al., 2006; Ito et al., 1998), our findings indicate widespread microglial activation in the *Spg15^-/-^* CNS. MotiQ analysis (Hansen et al., 2022), using Iba1 signal for high-throughput comparative morphology analyses, revealed that *Spg15^-/-^* microglia are less ramified in the dorsal thalamus (Fig. 1H), which is correlated with an activated state and provides evidence for microglial activation in a sensory brain region.

**Figure 1.**
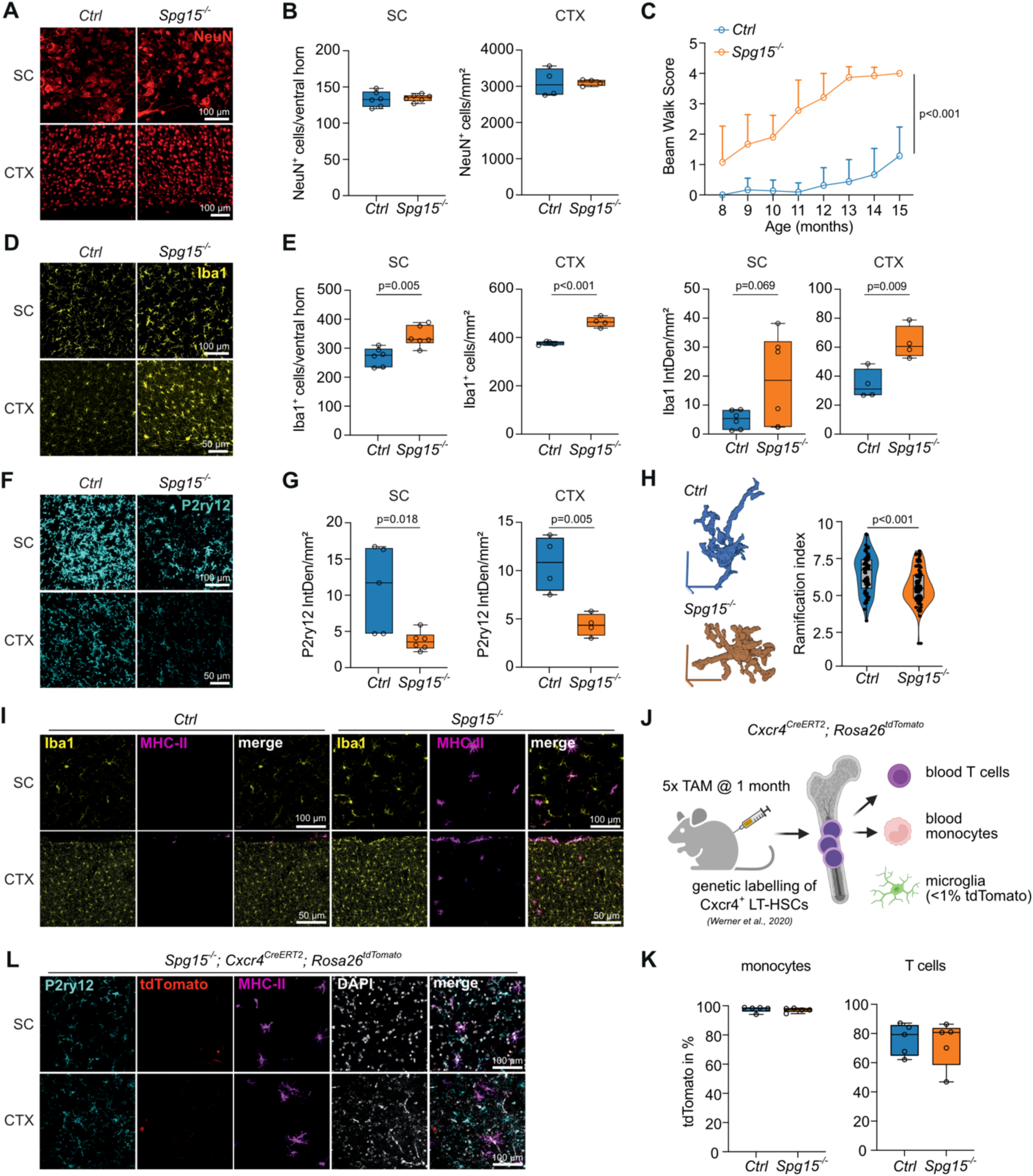
Microglial activation in the CNS of *Spg15^-/-^* mice. **A, B.** Representative immunofluorescent (IF) staining for NeuN (**A**) and quantification of NeuN^+^ neurons (**B**) in the ventral horn of the spinal cord (SC) as cells per ventral horn and layers V / VI of the primary motor cortex (CTX) as cells per mm^2^ in old (12-15 months) control *(Ctrl)* and *Spg15^-/-^* mice. **C.** Motor impairment of *Ctrl* and *Spg15^-/-^* mice at the indicated age as assessed by beam walk test (n = 7-23 per time-point and genotype). **D, E.** Representative IF staining for Iba1 (**D**) as well as quantification of Iba1^+^ cells per ventral horn (SC) or mm^2^ (CTX) and of Iba1 IF signal as integrated density (IntDen) per mm^2^ (**E**) in SC and CTX of old *Ctrl* and *Spg15^-/-^* mice. **F, G.** Representative IF staining for P2ry12 (**F**) and quantification of P2ry12 IF signal as IntDen per mm^2^ (**G**) in SC and CTX of old *Ctrl* and *Spg15^-/-^* mice. **H.** MotiQ morphometric analysis of the ramification of Iba1^+^ cells in the dorsal thalamus of old *Ctrl* and *Spg15^-/-^* mice (n = 60 cell from 3 mice for *Ctrl* and n = 95 from 4 mice for *Spg15^-/-^*). The 3D pictures show representative Iba1^+^ cells analyzed for their ramification index after reconstruction from confocal z stacks. **I.** Representative IF staining for MHC-II and Iba1 in SC and CTX from old *Ctrl* and *Spg15^-/-^* mice. **J.** Scheme of the *Cxcr4^CreERT2^; Rosa26^tdTomato^* fate-mapping model used in **K-L**. Mice were injected with tamoxifen (TAM) at four weeks of age, resulting in efficient labelling of long-term hematopoietic stem cells (LT-HSCs) and their progeny, but not microglia. Created with Biorender.com. **K.** Labeling efficiency of Ly6C^+^ blood monocytes and circulating CD3^+^ T cells in old *Spg15^+/+^; Cxcr4^CreERT2^; Rosa26^tdTomato^ (Ctrl)* and *Spg15^-/-^; Cxcr4^CreERT2^; Rosa26^tdTomato^* (*Spg15^-/-^*) mice. **L.** Representative IF staining for MHC-II, P2ry12, and tdTomato in SC and CTX from old *Spg15^-/-^; Cxcr4^CreERT2^; Rosa26^tdTomato^*mice. Statistical significance in **C, E, G, H**, and **N** was assessed with an unpaired Student’s *t*-test, for **C** only the p-value for the 15 months time point is shown.

Next, we evaluated MHC-II expression, which is often upregulated by activated microglia and/or recruited macrophages during neuroinflammation (Chen and Colonna, 2021). While, as expected, MHC-II was only detected in meningeal border-associated macrophages (BAM) in controls (Mrdjen et al., 2018), a subset of microglia showed a strong MHC-II signal in the *Spg15^-/-^* CNS parenchyma (shown for cortex and SC in Fig. 1I). To address whether the MHC-II expression originates from recruited monocyte-derived macrophages, we used *Cxcr4^CreERT2^; Rosa26^tdTomato^* fate-mapping (Fig. 1J) to detect HSC-derived cells (Werner et al., 2020). We crossed this fate-mapper with *Spg15^-/-^* (hereafter called *Spg15^-/-^*; *Cxcr4^CreERT2^; Rosa26^tdTomato^*) and control mice and induced tdTomato expression by tamoxifen injection at 4 weeks of age resulting in almost 100% labeling efficiency in monocytes in 12-15 month-old animals (Fig. 1K). Immunofluorescent staining of MHC-II, Iba1 and P2ry12 revealed that Iba1^+^ and P2ry12^+^ cells present in the CNS parenchyma were tdTomato-negative (shown for P2ry12 in Fig. 1L), indicating that MHC-II^+^Iba1^+^ and MHC-II^+^P2ry12^low^ cells were yolk sac-derived microglia and not of monocytic origin. In summary, our findings demonstrate that *Spg15^-/-^* mice develop widespread microglial activation before the onset of neuronal loss.

### Microglia acquire a DAM-like phenotype throughout the CNS of *Spg15^-/-^* mice

Next, we investigated the immune cell landscape in young (2-3 months) and old (12-15 months) *Spg15^-/-^*; *Cxcr4^CreERT2^; Rosa26^tdTomato^* mice and littermate controls to characterize a possible contribution of different resident and recruited immune cells to disease onset and progression. To this end, we used a UMAP representation of flow cytometry data comprising 855,105 CD45^+^ cells from different CNS regions (cerebral cortex, diencephalon, brain stem, cerebellum, and lumbar SC) of young and old control and *Spg15^-/-^* mice and overlaid it with the tdTomato signal (Fig. S1A, Fig. 2A). After categorizing the cells into T cells, neutrophils, BAM, and microglia based on their expression of cell-specific markers (Fig. 2B), we found that, as expected, almost all neutrophils exhibited tdTomato expression (Fig. 2A). While T cells consisted of tdTomato-positive and -negative cells, only few BAMs and virtually no microglia showed tdTomato positivity (Fig. 2A, C). We subsequently categorized the distinct CD45^+^ cells by age and genotype to determine their relative abundance in the respective condition (Fig. 2D). While the proportion of neutrophils and BAMs remained mostly constant with age, there was a notable decrease in the relative abundance of microglia in *Spg15^-/-^* mice, with a concomitant expansion of the T cell compartment during disease progression (Fig. 2D). Further assessment of the flow cytometry data showed no difference between control and *Spg15^-/-^* mice in terms of the relative contribution of different CNS regions to the total microglia within the CNS (Fig. 2E). This corresponds to our immunohistological findings showing microglia expansion in cortex and SC, suggesting that microglia expanded throughout the CNS in *Spg15^-/-^*mice.

**Figure 2.**
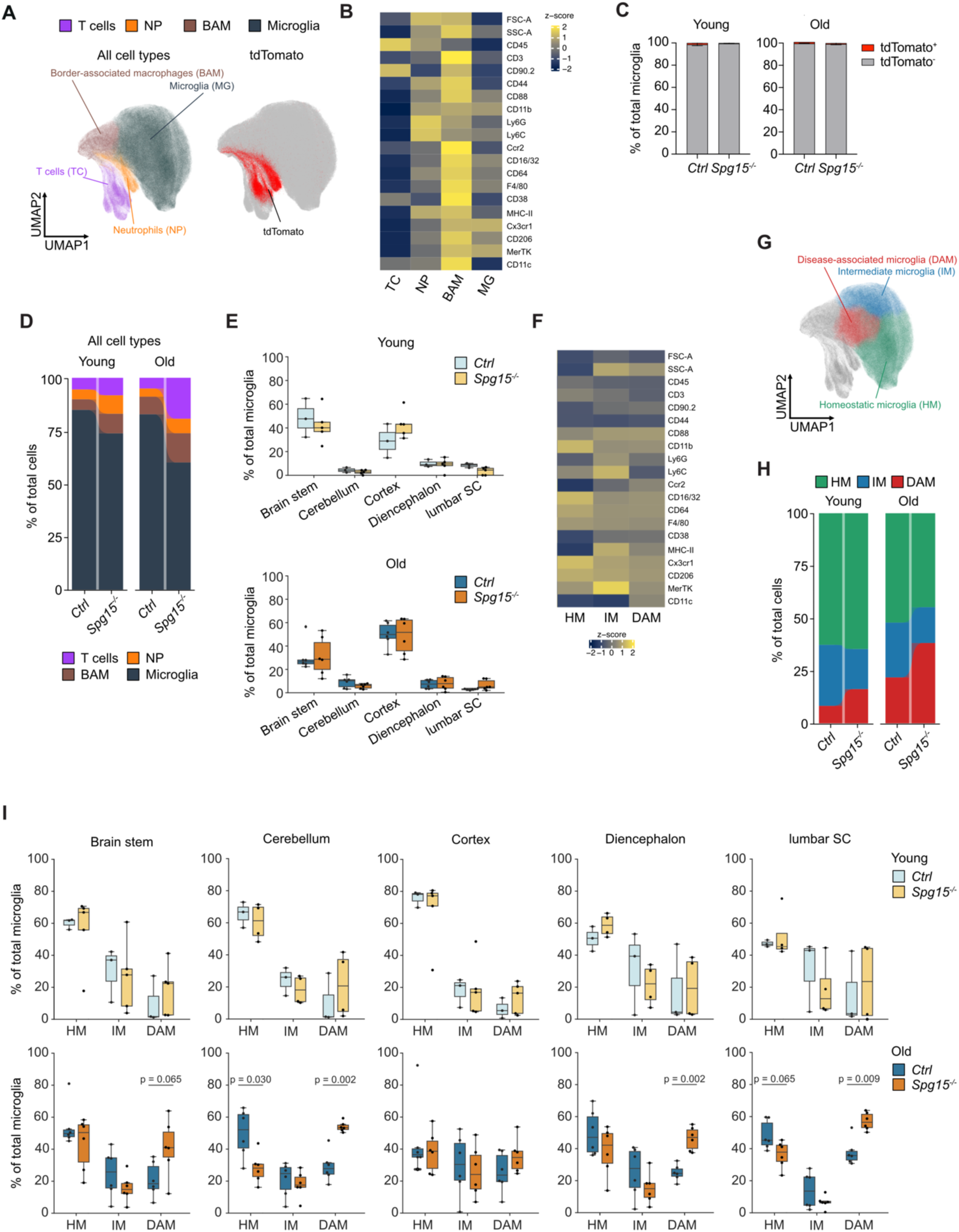
Characterization of the immune cell landscape in the CNS of young and old *Spg15^-/-^* mice. **A.** Left: Unsupervised Uniform Manifold Approximation and Projection (UMAP) plot of a total of 855,105 CD45^+^ cells from young and old *Spg15^+/+^; Cxcr4^CreERT2^; Rosa26^tdTomato^ (Ctrl)* and *Spg15^-/-^; Cxcr4^CreERT2^; Rosa26^tdTomato^* (*Spg15^-/-^*) mice, isolated from various regions across the central nervous system (CNS) including brain stem, cerebellum, cerebral cortex, diencephalon, and lumbar spinal cord (SC). Cells were manually assigned into four cell types or cell states based on their surface marker expression. Right: tdTomato^+^ cells are displayed on the UMAP plot including all CD45^+^ cells (n=3-6 per age and genotype). **B.** Pseudo-bulk heatmap of surface markers detected in flow cytometry analysis determining cell types and subtypes. **C.** tdTomato-labeling of microglia in young (left) and old (right) *Spg15^+/+^; Cxcr4^CreERT2^; Rosa26^tdTomato^ (Ctrl)* and *Spg15^-/-^; Cxcr4^CreERT2^; Rosa26^tdTomato^* (*Spg15^-/-^*) mice. **D.** Percentage of cell types or cell subtypes as defined in **A** of total CD45^+^ cells in either young or old mouse brains separated by genotypes. **E.** Contribution of different CNS regions (brain stem, cerebellum, cortex, diencephalon, lumbar SC) to relative microglia numbers in either young (upper panel) or old (lower panel) *Ctrl* or *Spg15^-/-^* mice. **F.** Pseudo-bulk heatmap of surface markers detected in flow cytometry for microglia subtypes. **G.** UMAP plot of microglia subtypes subset from plot in **A**. **H.** Percentage of microglia subtypes in all microglia in either young or old mouse CNS separated by genotypes. **I.** Percentage of microglia subtypes in either young (upper panel) or old (lower panel) animals across CNS regions. Statistical significance was assessed with unpaired t test with Welch correction. BAMs: border-associated macrophages; DAM: disease-associated microglia; HM: homeostatic microglia; IM; intermediate microglia; NP: neutrophils; TC: T cells.

As microglia may exist as distinct (activated) subpopulations in neuroinflammatory conditions, we used additional surface markers and cell size/granularity properties (Fig. S1A, Fig. 2F) to subcategorize microglia further. We found three microglial subpopulations across different CNS regions, which we termed as homeostatic microglia (HM) defined by high Cx3cr1 expression, intermediate microglia (IM) due to their increase in size (FSC-A), granularity (SSC-A) and MHC-II expression compared to HM, and DAM due to their MHC-II and CD11c expression (Fig. 2F, G, Fig. S1A). Relative quantification of all microglia subpopulations of young and old control and *Spg15^-/-^* mice revealed a slight expansion of DAM in 2-3 months young and substantial expansion of DAM in 12-15 months *Spg15*^-/-^ mice compared to age-matched controls (Fig. 2H).

To determine potential regional differences in the expansion of DAM, we assessed the relative abundance of HM, IM and DAM among all microglia in different CNS areas (Fig. 2I, Fig. S1B). This showed that DAM expanded significantly in diverse CNS regions of old but not young *Spg15^-/-^* mice compared to age-matched controls (Fig. 2I). Together, single-cell phenotyping of immune cell populations using surface markers points to conversion of HM into DAM throughout the CNS of old *Spg15^-/-^*mice before onset of neuronal loss, similar to observations in Alzheimer’s disease models (Chen and Colonna, 2021; Deczkowska et al., 2018).

### *Spg15^-/-^* microglia transition into a pro-inflammatory state

To characterize the functional state of microglia in old *Spg15^-/-^*; *Cxcr4^CreERT2^; Rosa26^tdTomato^* mice, we performed single-cell RNA-sequencing (scRNA-seq) of sorted CD11b^+^ cells (Fig. 3A) in combination with a proteogenomics approach (TotalSeq), which allowed combining transcriptome analysis with quantitative measurement of 44 surface proteins. In the UMAP representation of the transcriptome data, we identified three microglia subpopulations with *Apoe*^high^ *Lpl*^high^ *Spp1^high^* cells corresponding to DAM and *P2ry12^high^ Tmem119^high^ Csf1r^high^* cells to HM and IM (Fig. 3B). tdTomato transcripts were not detected in microglia but in neutrophils and BAMs as expected (Fig. S1C, D). In concordance with the flow cytometry data (Fig. 2), pseudotime analysis (Cao et al., 2019) confirmed that IM are likely a transitioning microglia population between HM and DAM (Fig. S1E).

**Figure 3.**
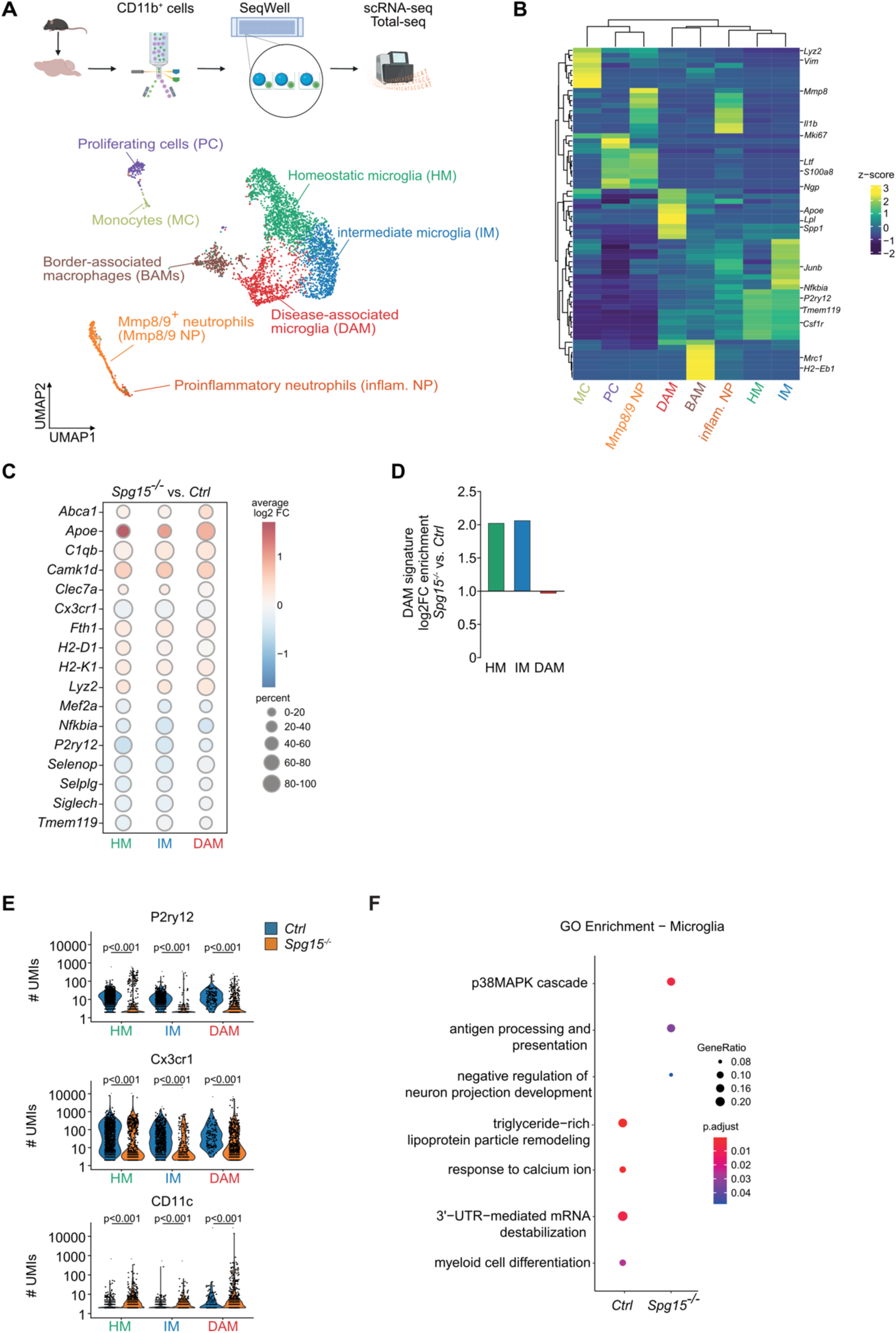
*Spg15^-/-^* microglia transition into damage-associated microglia (DAM). **A.** Top: Scheme representing the isolation procedure and single-cell RNA-sequencing (scRNA-seq) of CD11b^+^ cells (cells pooled from n = 2 mice per genotype). Created with Biorender. Bottom: UMAP plot showing 5,378 CD11b^+^ cells isolated by flow cytometry from old *Spg15^+/+^; Cxcr4^CreERT2^; Rosa26^tdTomato^ (Ctrl)* and *Spg15^-/-^; Cxcr4^CreERT2^; Rosa26^tdTomato^*(*Spg15^-/-^*) mice. Cells were manually assigned to eight cell types or cell states based on gene expression. **B.** Pseudo-bulk heatmap of Differentially Expressed Genes (DEGs) between clusters shown in **A**. DEGs were identified using the Wilcoxon rank-sum test. Resulting genes were filtered for a log_2_ fold change > 0.25 and genes expressed in at least 10% of cells in the respective clusters. Canonical markers used for annotation of clusters are highlighted. **C.** DEGs across microglia subpopulations comparing *Spg15^-/-^* to *Ctrl* brains (p.adjust<0.05). Log_2_ fold-change (log_2_FC) is indicated by red (upregulated in *Spg15^-/-^*) or blue (downregulated in *Spg15^-/-^*). **D.** Module score for a DAM gene signature was calculated for each cell in the microglia subclusters. log_2_FC were calculated, comparing *Spg15^-/-^* to the median of the *Ctrl*. **E.** Protein expression based on the number of Unique Molecular Identifiers (UMIs) aligned to the respective oligos in the TotalSeq comparing microglia from *Ctrl* to *Spg15^-/-^* mice. Statistical significance testing was performed using the Wilcoxon rank-sum test. **F.** Gene Ontology Enrichment Analysis (GOEA) performed on DEGs from **B.** Hypergeometric test with all genes in the “biological process” database as background, for statistical enrichment testing, in combination with the Benjamini-Hochberg procedure for multiple testing correction was performed. Ontology terms were filtered for a q-value > 0.2 and biological significance. BAMs: border-associated macrophages; DAM: disease-associated microglia; HM: homeostatic microglia; IM; intermediate microglia; NP: neutrophils; TC: T cells.

Comparison of differentially expressed genes (DEGs) in all three subpopulations between *Spg15^-/-^* mice and controls indicated that DAM from *Spg15^-/-^* mice express even higher levels of DAM-specific markers in comparison to littermate controls, such as *Apoe, C1qb*, and *Fth1* (Fig. 3C) (Mathys et al., 2017; Deczkowska et al., 2018; Chen and Colonna, 2021). However, also HM and IM populations from *Spg15^-/-^*mice upregulated DAM-specific genes (Fig. 3C). Therefore, we calculated a DAM signature score (see methods) indicating a significant increase of the expression of DAM-related genes in *Spg15^-/-^* HM and IM in comparison to controls (Fig. 3D, Fig. S1F). Further, we observed a downregulation of homeostatic microglial genes, such as *P2ry12*, *Tmem119,* and *Nfkbia* (Fig. 3C), which are also known to be downregulated in HM when transitioning into DAM (Chen and Colonna, 2021). This data suggested that HM and IM in *Spg15^-/-^* mice are starting to convert into DAM-like cells. We confirmed this observation using the quantitative TotalSeq approach, where we detected a significant decrease of the homeostatic markers P2ry12 and Cx3cr1 and upregulation of CD11c encoded by *Itgax* in all microglia populations from *Spg15^-/-^* mice compared to controls (Fig. 3E).

Next, we performed a Gene Ontology (GO) enrichment analysis of DEG from all three microglia populations comparing *Spg15^-/-^* with controls. While *Spg15^-/-^* microglia lost the expression of genes belonging to homeostatic terms, such as ‘*response to calcium ion*’ and ‘*triglyceride-rich lipoprotein particle remodeling*’, they upregulated genes within the terms ‘*antigen processing and presentation*‘, in line with the observed MHC-II expression, as well as ‘*p38MAPK cascade*’ indicating that these cells undergo cellular stress (Fig. 3F). Together, our multi-omics approach validated the observed DAM gene signature using flow cytometry and revealed a general conversion of *Spg15^-/-^*microglia subpopulation towards DAM-like phenotype.

### Expansion of CD8^+^ T cells with an effector-like phenotype in the CNS of *Spg15^-/^*^-^ mice before onset of neuronal loss

Antigen presentation in an MHC-dependent context is crucial for local activation of T cells in inflammatory CNS disorders (Berriat et al., 2023; Goverman, 2009). Given the increased MHC-II expression in microglia and the increase of T cells in *Spg15^-/-^* mice in flow cytometry (Fig. 1, Fig. 2), we examined potential T cell infiltration in more detail. Indeed, we observed an increased number of CD3^+^ T cells per CNS in 2-3 month-old young and 12-15 month-old *Spg15^-/-^* mice compared to age-matched controls (Fig. 4A). The general increase of CD3^+^ T cells was confirmed via immunofluorescent staining in the SC and cortex in old *Spg15^-/-^* mice (Fig. S2A, B). In the SC, CD3^+^ T cells were most abundant in white matter, which corresponds to the prominent microglial activation in the ventral horn of the lumbar SC (Fig. S2B). These findings were similar to the T cell infiltration observed in old *Spg11^-/-^* mice (Hörner et al., 2022). Interestingly, our fate-mapping model, where ∼80% of T cells in the blood are tdTomato-positive (Fig. 1K), revealed that only ∼50% of CNS-resident CD3^+^ T cells were positive for tdTomato (Fig. 4B). We confirmed the presence and equal distribution of tdTomato-positive and -negative CD3^+^ T cells via immunofluorescence in old *Spg15^-/-^*; *Cxcr4^CreERT2^; Rosa26^tdTomato^* mice (Fig. 4C, D), indicating that microglia activation and neurodegeneration caused by *Spg15* deficiency leads to a recruitment of T cells generated in the bone marrow after tamoxifen administration and a local expansion of T cells that have entered the brain in the first weeks of life.

**Figure 4.**
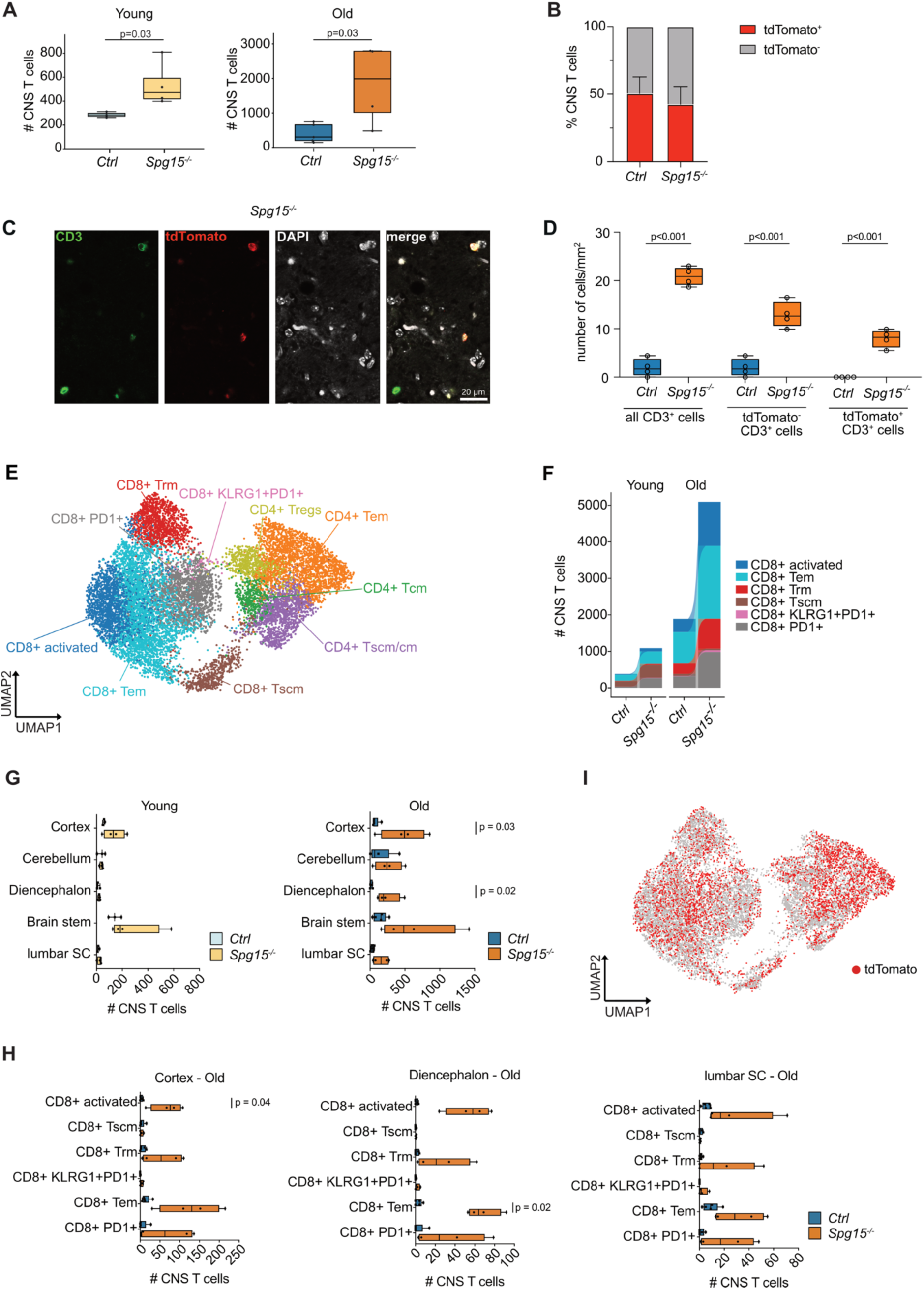
High-dimensional analysis of the T cell compartment in the CNS of *Spg15^-/^*^-^ mice. **A.** Number of CNS CD3^+^ T cells in young (2-3 months) and old (12-15 months) *Spg15^+/+^; Cxcr4^CreERT2^; Rosa26^tdTomato^ (Ctrl)* and *Spg15^-/-^; Cxcr4^CreERT2^; Rosa26^tdTomato^* (*Spg15^-/-^*) mice analyzed by flow cytometry. **B.** Flow cytometry analysis of tdTomato expression in CD3^+^ T cells present in the CNS of old *Ctrl* and *Spg15^-/-^*mice. Data is represented as mean with SEM. **C.** Representative IF double-staining for CD3 and tdTomato in the brain of 12-15 month-old *Spg15^-/-^; Cxcr4^CreERT2^; Rosa26^tdTomato^* mice. **D**. Quantification of CD3^+^ cells per mm^2^ in the CTX of 12-15 month-old *Ctrl* and *Spg15^-/-^* mice. **E.** Unsupervised UMAP of CD3*^+^* T cell clusters from control and *Spg15^-/-^* mouse brains. T cells were pooled from young and old animals and annotated according to key marker protein expression analyzed by flow cytometry. **F.** Compositional changes of CD8^+^ T cell subtypes as defined in **E** in young and old *Ctrl* and *Spg15*^-/-^ mice. **G.** Number of CD3^+^ T cells in different brain regions shown for young and old *Ctrl* and *Spg15*^-/-^ mice. **H.** Compositional changes of CD3^+^ T cell subsets as defined in **E** in cortex, diencephalon, and lumbar SC of old control and *Spg15^-/-^* mice. **I.** tdTomato expression overlaid on the UMAP. **A, D, G** Statistical significance was assessed with an unpaired Student’s *t*-test. **H** Statistical significance was assessed with an one-way FDR-corrected ANOVA.

Next, we characterized the pool of CD3^+^ T cells in more detail by flow cytometry and could observe that both the CD8^+^ and the CD4^+^ T cell compartments were expanded in old *Spg15^-/-^* mice with particularly CD8^+^ T cell numbers being strongly increased (Fig. S2C, D). To further subclassify CNS T cells, we clustered all T cells irrespective of the experimental group based on the expression of canonical surface markers (Figs. 4E, Fig. S2E). This resulted in a clear separation of CD4^+^ and CD8^+^ T cells, both of which could be further subdivided into phenotypically defined subsets. Specifically, there were six distinct subtypes of CD8^+^ T cells, namely CD62L^+^CD44^low^ stem-like central memory T cells (Tscm), CD44^+^CD69^+^CD62L^low^ effector memory T cells (Tem), CD44^high^CD25^+^CD69^+^ activated T cells (activated), tissue-resident CD69^+^CD103^+^ T cells (Trm), PD1^+^CD69^+^CD103^-^ effector T cells (PD1^+^) and PD1^+^ T cells co-expressing KLRG1 (KLRG1^+^PD1^+^). In the CD4 compartment, we detected four subtypes, which comprised CD4^+^CD25^high^ICOS^+^ Treg cells as well as CD62L^+^CD44^low^ stem-like central memory cells (Tscm), CD62L^low^CD44^+^ central memory cells (Tcm), and CD69^+/-^ effector memory cells (Tem) (Fig. 4E). Next, we assessed how these T cell subpopulations were distributed across the experimental groups (Fig. S2F). This showed a shift from a stem-like phenotype present in the CD8^+^ and CD4^+^ compartments in young *Spg15*^-/-^ and control mice towards an effector-like state in old animals, which was exacerbated in the *Spg15^-/-^* cohort (Fig. 4F, Fig. S2G, H). This overall increase in effector-like states was dominated by increases in the populations of activated, effector memory, tissue-resident, and PD1-expressing effector CD8^+^ T cells (Fig. 4F, Fig. S2G).

Based on the distinct neuropathology and expansion of DAM-like microglia in the five different CNS areas (Fig. 2I), we also assessed the expansion of T cells in these areas and observed an expansion of the total pool of CD3^+^ T cells in all areas with the highest enrichment in the cortex and diencephalon (Fig. 4G, Fig. S3A, B). Further subclassification of T cells in these areas indicated an expansion of activated, effector memory, tissue-resident, and PD1^+^ effector CD8^+^ T cells as detected for the complete CNS (Fig. 4H). This supports the notion that a potential CD8^+^ T cell contribution to CNS pathology is not restricted to an individual area but present throughout the CNS reflective of the microglial activation while the response within the CD4^+^ T cell compartment was more equivocal (Fig. S3C-E).

Next, exploiting the strength of the *Cxcr4^CreERT2^* fate-mapping model that allows the discrimination of locally proliferating versus recruited T cells into the CNS, we quantified the tdTomato-positive fraction of T cells. The tdTomato signal was detected in all T cell clusters (Fig. 4I), suggesting that T cell differentiation was independent of the time point of entry into the CNS. Analysis of all six CD8^+^ and four CD4^+^ T cell subpopulations in old *Spg15*^-/-^ mice and littermate controls revealed a distinct behavior of these different T cell types. While there was not a big difference in the CD4^+^ compartment across genotypes, we detected a slight increase of tdTomato-positive CD8^+^KLRG1^+^PD1^+^ T cells in *Spg15*^-/-^ mice compared to controls, indicating either an increased recruitment of cells into the CNS or preferential local proliferation of tdTomato-positive cells (Fig. S3F). In contrast, we observed a substantial enrichment of tdTomato-negative CD8^+^ Trm and Tscm compartments (Fig. S3F), supporting the idea that particularly CD8^+^ T cells can be recruited into the CNS before tamoxifen treatment where they locally expand over time.

In summary, these data support the notion that the T cell compartment, and mainly CD8^+^ T cells with a tissue-resident and effector phenotype, are expanded in *Spg15*^-/-^ mice. This expansion is already detectable at early age but more prominent in old animals and involves both early-infiltrating tdTomato-negative and late-infiltrating tdTomato-positive CD8^+^ T cells. Like the phenotype shift of microglia, T cell activation was not restricted to motor regions and might thus contribute to the complicated neuropathology of SPG15 involving motor, sensory, and cognitive impairment (Pensato et al., 2014).

### scRNA-seq of T cells in *Spg15^-/-^* mice identifies a type-I IFN/PD1 coexpressing subpopulation of CD8^+^ T cells

To better define the differentiation states of T cells in the brain of old *Spg15^-/-^* mice, we performed 384-well-based cell sorting of CD3^+^ cells with subsequent transcriptomic analysis using Smart-Seq2 (Picelli et al., 2014) (Fig. S4A). After data quality checks and filtering for pure αβ T cells, this resulted in a dataset of 526 high-quality single cells. In the UMAP plot (Fig. 5A, Fig. S4B), four distinct CD8^+^ T cell clusters were identified, which largely corresponded to four of the clusters of effector-like CD8^+^ T cells previously identified by flow cytometry (Fig. 4E). Presence of only 61 cells in the CD4^+^ T cell compartment prevented further subtyping (Fig. 5A, Fig. S4B). By plotting marker genes for the respective T cell clusters including genes that differentiate between effector CD8^+^ T cell states, we identified a clearly separated PD1-expressing type-I IFN signaling (IFN-I)/PD1 CD8^+^ T cell cluster (*Tbx21*, *Ly6c2*, *Ifit3, Gzmb, Klrg1*) (Hussain and Quinn, 2019; Zöphel et al., 2022; Istaces et al., 2019) corresponding to the PD1^+^KLRG1^+^ CD8^+^ T cell cluster we detected by flow cytometry (Fig. 4E). Furthermore, we identified a PD1^+^ effector state of CD8^+^ T cells (CD8^+^PD1^+^), characterized by high expression of multiple inhibitory receptors (*Pdcd1*, *Ctla4*, *Lag3*, *Tigit, Havcr2*/TIM-3) and *Tox* (E Hirbec et al., 2017; Wherry and Kurachi, 2015), a cluster of activated/effector memory CD8^+^ T cells (Tact/em) co-expressing *Itgax*, *Eomes, Klre1, and Cd69* (Shinoda et al., 2012; Chen et al., 2023), a CD8^+^ tissue-resident (Trm) cluster with co-expression of *Itgae, Cd69, Rorgt*-associated genes (*Stat3, Il17f*), *Ifng* and *Bhlhe40* (Hussain and Quinn, 2019; Zöphel et al., 2022; Istaces et al., 2019), and a small CD4^+^ T cell cluster.

**Figure 5.**
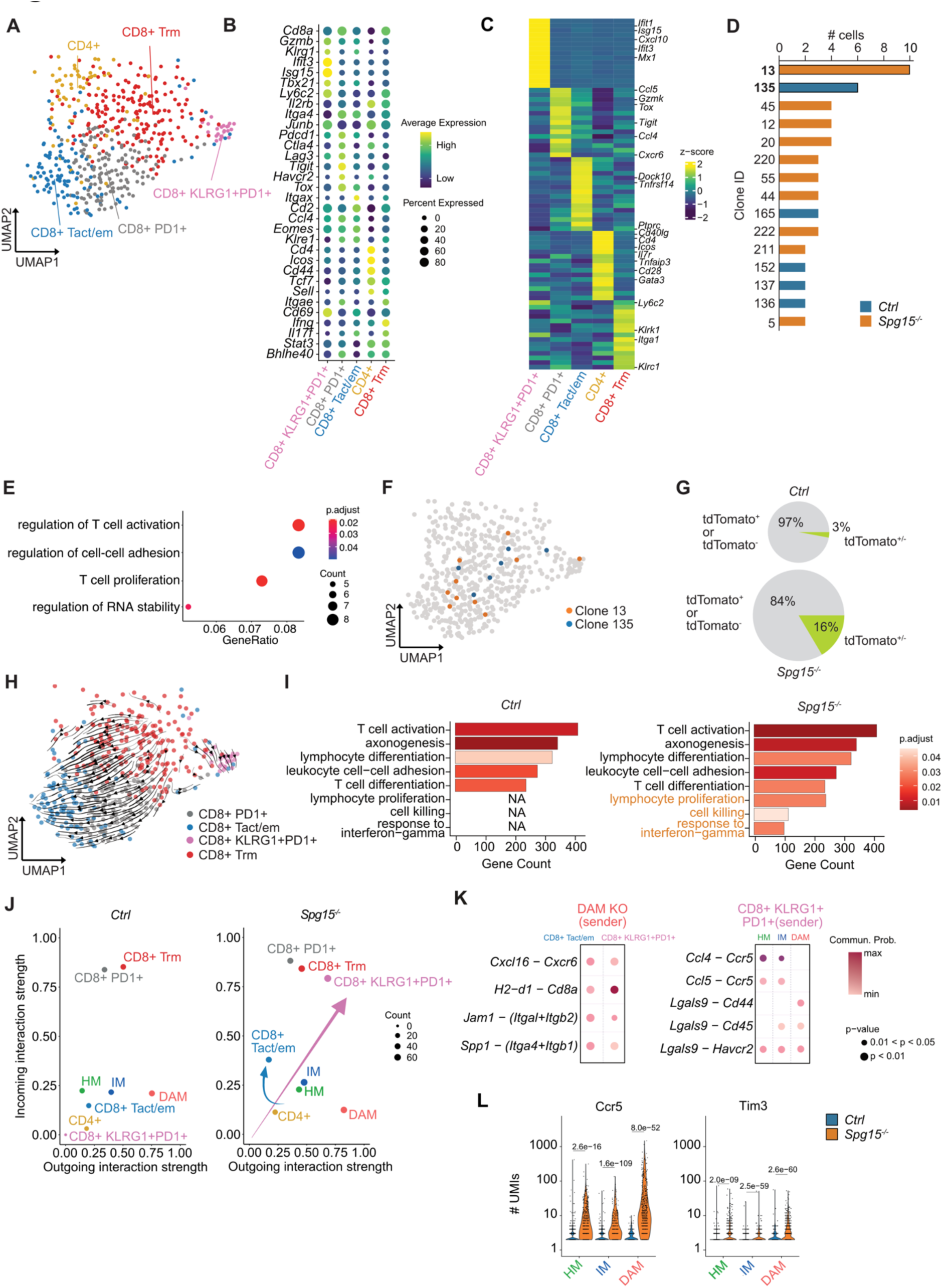
scRNA-seq analysis reveals intricate bidirectional communication between DAM and CD8^+^ T cells in the CNS of *Spg15^-/-^* mice. **A.** UMAP from full length scRNA-seq data generated with SmartSeq2 from CNS T cells isolated from the CNS of 12-15 month-old *Ctrl* and *Spg15^-/-^* mice. **B.** Dotplot visualizing marker genes used for annotation of the clusters in panel **A**. Genes were selected based on the high expression and previous association with CD8^+^ T cell differentiation. **C.** Pseudo-bulk heatmap showing upregulated DEGs identified between cell clusters. DEGs were identified using the Wilcoxon rank-sum test. Resulting genes were filtered for a log_2_ fold change > 0.1 and genes expressed in at least 30% of cells in the respective clusters. Genes that were used to further characterize the cell clusters were highlighted. **D.** T cell clones inferred with TRUST4 were quantified with Scirpy. Dominant clones in *Ctrl* (135, blue) or *Spg15^-/-^* (13, orange) are highlighted. **E.** GO terms obtained with GO enrichment analysis performed with genes differentially expressed between the dominant clones in *Spg15^-/-^*(13) and *Ctrl* mice (135). **F.** Visualization of the single cells from the two dominant clones in *Ctrl* (135, blue) and *Spg15^-/-^*animals (13, orange) in the UMAP from panel **A**. **G.** Percentage of either tdTomato^+^ or tdTomato^-^ CD8^+^ T cell clones (grey) as well as CD8^+^ T cell clones containing both tdTomato^+^ and tdTomato^-^ cells (green) in old *Ctrl* or *Spg15^-/-^* animals. The size of the pies represents the relative number of cells in the respective condition. **H.** Averaged velocity vectors, representing the direction and strength of development based on the ratios of exons and introns, were calculated on CD8^+^ T cells only and plotted onto the UMAP from panel A. **I.** DEGs between CD8^+^ T cells from *Ctrl* and *Spg15^-/-^* animals were used to perform GO enrichment analysis and enriched GO terms plotted. Terms uniquely enriched in *Spg15^-/-^*animals were highlighted in orange. **J.** Inference of cell-cell interaction using CellChat contrasting incoming (y-axis) and outgoing interaction strengths (x-axis, arbitrary unit) visualized for microglia and T cells from mouse brains from *Ctrl* (left) and *Spg15^-/-^* animals (right). The number of interactions was coded as the dot size of the cell cluster. Clusters with drastic changes in communication behavior were highlighted with arrows in the *Spg15^-/-^* (right) plot. **K.** Ligand-receptor pairs between (left) DAM (sender) and CD8^+^ Tem or KLRG1^+^PD1^+^ CD8^+^ T cells (receiver) and (right) KLRG1^+^PD1^+^ CD8^+^ T cells as sender with HM, IM, and DAM subtypes as receivers. Communication probability was color-coded on a continuous scale. **L.** Ccr5 and Tim3 expression represented as the number of UMIs derived from TotalSeq data was visualized for microglia subsets. Statistical significance was assessed using a Wilcoxon rank sum test.

Differential gene expression analysis identified subset-specific transcriptional patterns (Fig. 5C). Specifically, CD8^+^ Trm expressed high levels of *Itga1, Ly6c2*, *and Klrc1,* indicating long term memory, tissue residency, repeated stimulation, and cell division (Mackay et al., 2013; Siracusa et al., 2019; Cerwenka et al., 1998; Walunas et al., 1995; Borst et al., 2022). Increased activation of CD8^+^ Tact/em was indicated by expression of canonical activation/effector genes and higher expression of *Ptprc* and *Tnfrsf14*. We further noted expression of activation/effector markers, PD1-associated markers, and *Tox* in PD1^+^ CD8^+^ T cells, as well as a type I IFN signature in the IFN-I/PD1 CD8^+^ T cell subset (Fig. 5C). Based on previous characterization of the functional properties of these IFN-I/PD1 CD8^+^ T cells (Schetters et al., 2017; Chen et al., 2023; Ritzel et al., 2016) it is tempting to speculate that this subset of CD8^+^ T cells can actively support microglia activation. Mapping the expression of tdTomato onto the UMAP showed a homogeneous distribution of tdTomato over the T cell clusters suggesting that the time-point of entry into the CNS of CD8^+^ T cells does not influence their differentiation state (Fig. S4C).

Taken together, these data support the notion that effector-like CD8^+^ T cells expand in the CNS of *Spg15^-/-^* mice, extending the previously reported expansion of CD8^+^ T cells in *Spg11*-deficient animals (Hörner et al., 2022). As this accumulation of CD8^+^ T cells is also driven by a local expansion of T cells recruited early during the disease course, it is conceivable that the CD8^+^ T cell pool, both recruited early and late, is actively contributing to the development of neuroinflammation and subsequent neurodegeneration in *Spg15*^-/-^ mice.

### Clonal expansion of effector CD8^+^ T cells in *Spg15^-/-^*mice is independent of their transcriptional state and time-point of recruitment

Given that CD8^+^ T cells actively contribute to neuroinflammation in SPGs as evidenced for *Spg11* (Hörner et al., 2022) and suggested by our own findings in *Spg15*^-/-^ *mice*, we next asked if a clonal expansion can be detected in the CD8^+^ CNS T cell compartment in *Spg15^-/-^* mice. We used TRUST4 for the reconstruction of the T cell receptor repertoire from full-length Smart-Seq2 data (Song et al., 2021). This revealed more T cell clones with >3 cells in *Spg15*^-/-^ than in controls (Fig. 5D, Fig. S4D, E), with both genotypes exhibiting one dominant clone. To assess if the most prevalent clones of *Spg15^-/-^* mice and controls showed transcriptional differences, we assessed differential gene expression. This identified genes associated with T cell effector function, proliferation or activation, such as *Klrc2, Jak1,* and *Eif5* (Tan et al., 2022; Ross et al., 2016; Li et al., 2022) in the *Spg15*^-/-^ clone (Fig. S4F), suggesting a functionally more active phenotype of this expanded CD8^+^ T cell clone. The shift of *Spg15^-/-^* clones into a more activated state was confirmed by gene set enrichment analysis (GSEA) of the top 100 DEG between the *Spg15^-/-^* and control clones. Terms enriched for CD8^+^ T cells from *Spg15^-/-^* animals were associated with changes in T cell activation, adhesion, and proliferation (Fig. 5E). To answer if clonally expanded CD8^+^ T cells have defined T cell states, we mapped the expanded clones (Fig. S4G) and the two most prevalent clones (Fig. 5F) onto the UMAP. We did not detect enrichment of clones in a specific CD8^+^ T cell subset (Fig. S4G). This suggests that the TCRs of the clonally expanded CD8^+^ T cells are not indicative of their transcriptional state but that this process is driven by alternative differentiation cues. This is further supported by the fact that within expanded CD8^+^ T cell clones in *Spg15*^-/-^ animals tdTomato^+^ and tdTomato^-^ CD8^+^ T cells can be detected at the same time (Fig. 5G, Fig. S4C). Beyond that, the clonal analysis suggested that CD8^+^ T cell clones from *Spg15*^-/-^ mice were enriched in effector subtypes while clones in controls were mainly Trm (Fig. S4H). In summary, these data support that clonal expansion of effector CD8^+^ T cells occurs in the *Spg15^-/-^* CNS. Based on their transcriptional state, the expanded clones might propagate a neuroinflammatory cascade in *Spg15^-/-^* mice.

### Increased intercellular communication between microglia and T cells in *Spg15*^-/-^ mice

Following the idea of a TCR-independent transcriptional differentiation of CD8^+^ T cells, we performed trajectory analysis and observed two separate pathways of differentiation. One from the pool of Trm over recently activated PD1^+^ T cells towards CD8^+^ Tact/em cells and the other from Trm cells into *Ly6c2* and type-I IFN-response genes expressing IFN-I/PD1 CD8^+^ T cells (Fig. 5H, Fig. S4I). As the path from Trm to CD8^+^ Tact/em cells was observed in both *Spg15*^-/-^ and controls, we aimed to uncover functional programs that were active across the different T cell states but enriched in CD8^+^ T cells isolated from *Spg15*^-/-^ animals. To this end, we performed GO term enrichment using a ranked gene list based on expression levels in the respective condition. This analysis revealed a T cell activation/differentiation pattern already present in CD8^+^ T cells isolated from controls, which was also detectable in cells from *Spg15*^-/-^ mice (Fig. 5I). Beyond this we identified additional enriched GO terms in CD8^+^ T cells from *Spg15^-/-^* mice like “*response to IFN-gamma*”, “*lymphocyte proliferation*”, and “*cell killing*” supporting that those cells were more activated than CD8^+^ T cells from controls (Fig. 5I). Next, we were interested in whether early recruitment into the CNS would affect functional properties of CD8^+^ T cells and analyzed tdTomato^+^ and tdTomato^-^ CD8^+^ T cells separately. Independent of tdTomato expression, CD8^+^ T cells from *Spg15^-/-^* mice expressed more genes associated with IFN-gamma production and positive regulation of neuron differentiation (Fig. S4J), suggesting that CD8^+^ T cells exert an active role in intercellular communication in the CNS of *Spg15*^-/-^ mice that is independent of the time of CNS entry (Fig. S4J).

This observation led us to analyze intercellular communication networks between T cells and microglia subsets using CellChat (Jin et al., 2021). To detect differences between cells from controls and *Spg15*^-/-^ mice, we inferred intercellular communication networks separately and subsequently jointly mapped them onto a shared two-dimensional manifold (Fig. 5J). This analysis revealed that microglia exhibited increased communication strength while transitioning from HM via IM to DAM. This supports our hypothesis that DAM act as senders towards CD8^+^ T cells. Here, Trm and PD1 CD8^+^ T cells are the main signal-receiving populations in *Ctrl* and *Spg15^-/-^* mice. In addition, CD8^+^ Tact/em cells and IFN-I/PD1 CD8^+^ T cells acted as strong receivers for signals emanating from IFN-I/PD1 CD8^+^ T cells in *Spg15^-/-^* mice.

The signaling from DAM to Trm and PD1 CD8^+^ T cells was enriched in four key immune cell signaling pathways (Fig. 5K, Fig. S4K), including *Cxcl16* - *Cxcr6*, MHC class I (*H2-d1*) - TCR (*Cd8a*), *Jam1* - Integrin α1β2, and Osteopontin (*Spp1*) - Integrin α4β1 signaling, suggesting that DAM-derived signals could contribute to the activation of CD8^+^ T cells in *Spg15*^-/-^ mice. A contribution of IFN-I/PD1 CD8^+^ T cells to the differentiation of HM towards DAM in *Spg15^-/-^* mice is supported by the notion that three communication pathways are directed from these T cells towards microglia, including chemokines (*Ccl4* - *Ccr4* and *Ccl5* - *Ccr5*) and galectins (Galectin 9 (*Lgals9*) - Tim3 (*Havcr2*)/*Cd45*/*Cd44*). To confirm that these proteins are expressed by microglia, we reassessed our TotalSeq data. This showed significantly higher expression of Ccr5 and Tim3 in all three microglia subsets in *Spg15^-/-^* mice compared to controls (Fig. 5L). These results reveal a potential communication network between microglia and CD8^+^ T cells as the basis for a neuroinflammatory state in the CNS of *Spg15^-/-^* mice.

## Concluding remarks

Immunomodulatory and genetic interventions can dampen disease progression in *Spg11^-/-^* mice, suggesting neuroinflammation as a driving factor in complicated HSP (Hörner et al., 2022). Using *Spg15^-/-^* mice, we provide the first comprehensive immunophenotyping in a model of complicated HSP. Our findings show that, unlike brain injuries and inflammatory CNS disorders, *Spg15*^-/-^ is not connected to infiltration of peripheral myeloid cells. Instead, microglia lose homeostatic functions and assume a DAM-like state throughout the CNS, which is associated with local expansion of early-infiltrating T cells, recruitment of late-generated T cells, as well as prominent activation of CD8^+^ T cells. Thus, widespread neuroinflammation driven by microglia and CD8^+^ T cells might underlie sensory, mental, and motor impairment in complicated HSP. We further reveal a potential communication network between microglia and CD8^+^ T cells for targeted anti-inflammatory interventions aiming to halt disease progression in complicated HSP.

## Acknowledgements

We thank Michael Kraut, Heidi Theis, Stephanie Weber and Svenja Bourry for technical assistance.

## Author contributions

Conceptualization, R.S., E.M., M.B.; Data acquisition, analysis, and interpretation, A.F., H.H., D.S., R.S., M.K., N.B.-S., C.O.-S., T.E., M.Bü., M.Bec., L.B., K.H.; Resources and discussion, C.A.H.; Supervision and conceptual discussion, R.S., E.M., M.B., Writing - original draft, A.F., H.H., R.S., E.M., M.B.; Editing, all authors.

## Material and Methods

### Animals

Mouse husbandry was in accordance with institutional and EU or national guidelines for animal use, approved by the competent authority (Thüringer Landesamt für Verbraucherschutz, TLV), and supervised by the institutional veterinarians. Animal procedures were performed in adherence to our project licenses issued by the federal state Thueringen (TLV administrative authorization number 02-056/16). Young animals were 2-3 month-old, old animals were between 12 and 15 months of age. All experimental and breeder mice were on a the C57BL/6 background and contained *Cxcr4^CreERT2x^*and *Rosa26^tdTomato^* (Ai14) alleles (Werner et al., 2020). These mice were crossed with *Zfyve26*^-/-^ or *Zfyve26*^+/-^ mice (Khundadze et al., 2013) to obtain *Spg15*^-/-^ (*Zfyve26*^-/-^) and control mice (*Zfyve26*^+/-^; *Zfyve26*^+/+^). Fluorescent-labeling of hematopoietic stem cells was performed by five sequential intraperitoneal injections of 1 mg tamoxifen per day at the age of four weeks (1 mg per day on five sequential days). Tamoxifen was prepared 1:10 (w/v) in ethanol (Carl Roth, 5054) and adjusted to 10 mg ml^−1^ in corn oil (Caelo, 7284).

### Beam walk

Motor deficits were assessed using a beam walk test. The beam was 1 m in length and 7 mm broad, with a platform at each end. The beam was mounted 20 cm above the surface of a table. At the far end of the beam, a cage with nesting material from the home cage was placed to attract the animal. Each animal had to cross the beam three times per session. Animals were trained for beam walk starting at the age of 8 months. Motor deficits were assessed once a month. All trials were video recorded for detailed analysis. The following scoring system was used for assessment: score 0, animal is sure-footed, hind legs are under the body when walking, tail swings from left to right; score 1, like score 0 but the tail is occasionally nestled on the beam; score 2, the tail is nestled on the beam and shows little movement, hind legs are still under the body, belly is lifted off the beam; score 3, signs like score 4, but not continuously; score 4: tail is nestled on the beam and shows little movement, animal continuously slides with belly on the beam and thus makes a hump, hind legs (one or both) are placed sideways on the beam; 5, like score 4 but also forelegs are placed sideways on the beam.

### Immunohistology, microscopy, and image analysis

Mice were killed by 5% isoflurane and then transcardially perfused with PBS followed by 4 % formaldehyde/PBS (pH 7.4). Cryosections (40 μm) were cut after cryoprotecting the tissues in 10% and 30% sucrose/PBS. Sections were stored and processed in a 10 mmol/l Tris, 10 mmol/l phosphate, 155 mmol/l NaCl working buffer (WB) adjusted to pH 7.4. Successive 30 min incubations with 50% methanol containing 0.3% H_2_O_2_ and 5% BSA containing 0.3% Triton-X100 were carried out in WB before applying primary antibodies for 24–72 h and fluorescent secondary antibodies for 2 h in in WB containing 1 % BSA and 0.3 % Triton-X100. Images were captured with the LSM 900 using ZEN software (Carl Zeiss Jena). Images were processed using ImageJ (v1.51) and Adobe Photoshop (v21.1.3). The following primary antibodies were used: anti-NeuN (Abcam #177487), 1:200; anti-CD3 (Abcam #56313), 1:5000; anti-P2ry12 (Anaspec AS-55053A), 1:500; anti-MHC-II (Biolegend #107622), 1:5000; anti-Iba1 (Abcam #5076), 1:400. Signal amplification using biotinylated secondary antibodies and fluorescent streptavidin was used to detect MHC-II and CD3. Cell counts and measurement of immunofluorescent signal intensities were performed on confocal images imported into ImageJ. NeuN^+^ cells in the ventral horn of the lumbar spinal cord were counted in a 500 µm x 500 µm region of interest (ROI) unilaterally placed immediately ventrolateral to the central canal, resulting in NeuN^+^ cells per ventral horn. NeuN^+^ and Iba1^+^ cells in the deep layers of the primary motor cortex were counted in 638 µm x 638 µm ROIs placed at the corpus callosum/layer VI boundary. Iba1 and P2ry12 signal intensities in the cortex were measured as Integrated Density (IntDen) in a similar ROI and then divided by the area. Iba1 and P2ry12 signal intensities in the ventral horn of the SC were measured as IntDen by selecting the white and grey matter ventrolateral to the central canal as ROI and then divided by the area. CD3 cells were counted in the deep layers of the primary motor cortex and the ventral spinal cord (grey and white matter) in 1 mm^2^ and 0.7 mm^2^ ROIs, respectively. MotiQ (Hansen et al., 2022) was used for morphometric analysis of Iba1^+^ microglia based on a customized Fiji plug-in. MotiQ cropper and thresholder were used in version v0.1.2 by using Huang for the intensity threshold. MotiQ 3D Analyzer version v0.1.5. was used with a minimum particle volume of 50 voxels. The ramification index is unit-free and was calculated as the ratio between the surface area of the cell and that of a sphere with the same volume and serves as a measure of the complexity of a cellular shape.

### Antibodies and flow cytometry analysis

Fluorescent-dye-conjugated antibodies were purchased from Becton Dickinson, BioLegend or eBioscience. Cell surface staining was performed at 4°C for 20 min with the addition of FcR-blocking reagents (1:100 CD16/32, 2% rat serum). Intracellular staining was conducted using the Foxp3 Staining Buffer Kit (eBioscience) with the addition of FcR-blocking reagents. Data were acquired on a BD LSRII or BD Symphony A5 flow cytometer (Becton Dickinson) and analyzed using FlowJo software package (FlowJo, LLC, v10.8.0), Cytoflow v.1.2, Pytometry v.0.0.1 (Buttner et al., 2022), Scanpy v.1.9.1 (Wolf et al., 2018) and scCoda v.0.1.4 (Büttner et al., 2021). Briefly, compensation was performed and cells were gated for CD45^+^ or CD3^+^ cells in FlowJo, respectively. The compensated fluorescence values for CD45^+^ or CD3^+^ gated cells were exported as .fcs files. These files were imported as Anndata (v0.8.0) objects for high dimensional analysis with Scanpy. After PCA, UMAP and Leiden clustering, cell clusters were annotated based on expression of canonical protein markers. Differential abundance testing was performed using scCODA. Statistical significance was assessed using the Mann-Whitney U test. For confirmation, manual gating for myeloid and T cell subsets was performed with Cytoflow and compared to the cell clusters generated with Scanpy.

### Purification of CNS immune cells

For isolation of immune cells, brains were separated in cortex, cerebellum, diencephalon, brainstem and spinal cord and minced in separate tubes. Subsequently, tissue was cut into small pieces, and incubated in a digestion mix (PBS containing 1 mg/ml collagenase D (Roche), 100 U/ml DNase I (Sigma-Aldrich), 2.4 mg/ ml of dispase (Gibco) and 3% fetal calf serum (FCS) (Invitrogen)) for 30 min at 37°C before mechanical disruption through a 100 μm filter on ice. Cells were then enriched using a one layer 27% Percoll density gradient centrifugation at 600 g for 5 min RT and the deceleration set to 3 out of 9. Peripheral blood was incubated with a lysis buffer (6.5 mM NH_4_C, 0.1 M KHCO_3_, 0.1 mM EDTA, pH 7.4) to lyse contaminating erythrocytes. Cells from peripheral blood were analyzed with BD Symphony A5. CD45^+^ brain cells were directly sorted on a BD Aria III or BD Symphony S6 cell sorter for sequencing experiments. The left hemisphere was used to sort CD45^+^ cells to perform scRNA-seq with SeqWell S^3 (Aicher et al., 2019). The right hemisphere was used for index sorting of CD3^+^ cells into 384-well plates (Penter et al., 2018) for scRNAseq with SmartSeq2 (Picelli et al., 2014). Only cells isolated with a purity of > 98% were used for further analysis.

### SeqWell S^3 sequencing library preparation for CNS CD45^+^ immune cells

CNS CD45^+^ cells from the brains of C57BL/6 *Spg15*^-/-^; *Cxcr4^CreERT2^; Rosa26^tdTomato^* or *Spg15*^+/-^;*Cxcr4^CreERT2^; Rosa26^tdTomato^* animals were isolated as described above. These cells were labeled with ADT antibodies (Biolegend) according to the manufacturer’s protocol for TotalSeq-A. 50 μl cell suspension with 1x10^6^ cells were resuspended in staining buffer (2% BSA, Jackson Immuno Research; 0.01% Tween-20, Sigma-Aldrich; 1x DPBS, GIBCO) and 5 μl mouse TruStain FcX FcBlocking (Biolegend) reagent were added. The blocking was performed for 10 min at 4°C. In the next step 1 μg unique TotalSeq-A antibody was added to each sample and incubated for 30 min at 4°C. After the incubation time 1.5 ml staining buffer were added and centrifuged for 5 min at 350 *g* and 4°C. Washing was repeated 3 times. Subsequently, the cells were resuspended in an appropriate volume of 1x DPBS (GIBCO), passed through a 40 μm mesh (Flowmi Cell Strainer, Merck) and counted, using a Neubauer Hemocytometer (Marienfeld). Cell counts were adjusted and hashtagged cells were pooled equally. The cell suspension was loaded with 25,000 cells to 1.1x10^5^ beads per SeqWell array. Reverse transcription, cDNA amplification and library generation were performed according to the recommendations from Aicher et al. (Aicher et al., 2019). The reads were aligned using STAR v.2.6.1a_08-27 (Dobin et al., 2013) to the mm10 mouse genome (Release M16 Gencode; GRCm38.p5). Raw data were uploaded to GEO under GSE244476.

### SmartSeq2 sequencing library preparation for CNS CD3^+^ T cells

CNS T cells from the brains of C57BL/6 *Spg15*^-/-^; *Cxcr4^CreERT2^; Rosa26^tdTomato^* or *Spg15*^+/-^;*Cxcr4^CreERT2^; Rosa26^tdTomato^* animals were isolated as described above. Cells were FACS sorted into eight 384-well plates containing 2.3 µl lysis buffer (Guanidine Hydrochloride (50 mM; Sigma–Aldrich: G3272), dNTPs (17.4 mM; NEB: N0447), SMART dT30VN primer (2.2 µM; IDT) retaining protein expression information for every well to subsequently match with the respective single-cell transcriptomic data in an index sorting approach. Plates were sealed and stored at −80 °C until further processing. Smart-Seq2 libraries were finally generated on a Tecan Freedom EVO and Nanodrop II (BioNex) system as previously described (Picelli et al., 2013). In short, lysed cells were incubated at 95 °C for 3 min. 2.7 µl RT mix containing SuperScript II buffer (Invitrogen: 18064071), 9.3 mM DTT, 370 mM Betaine (Sigma–Aldrich: B0300), 15 mM MgCl2 (Sigma–Aldrich: 63069), 9.3 U SuperScript II RT (Invitrogen: 18064071), 1.85 U recombinant RNase Inhibitor (Takara: 2313 A), and 1.85 µM template-switching oligo (Eurogentec) was aliquoted to each lysed cell using a Nanodrop II liquid handling system (BioNex) and incubating at 42 °C for 90 min and 70 °C for 15 min. 7.5 µl preamplification mix containing KAPA HiFi HotStart ReadyMix (KAPA: 7958935001) and 2 µM ISPCR primers (IDT) was added to each well and full-length cDNA was amplified for 16 cycles. cDNA was purified with 1× Agencourt AMPure XP beads (Beckman Coulter: A63882) and eluted in 14 µl nuclease-free water (Invitrogen: 15667708). Concentration and cDNA size was checked for select representative wells using a High Sensitivity DNA5000 assay for the Tapestation 4200 (Agilent: 5067-5592). cDNA was diluted to an average of 200 pg/µl and 100 pg cDNA from each cell was tagmented by adding 1 µl TD and 0.5 µl ATM from a Nextera XT DNA Library Preparation Kit (Illumina: FC-131-1096) to 0.5 µl diluted cDNA in each well of a fresh 384-well plate. The tagmentation reaction was performed at 55 °C for 8 min before removal of the Tn5 from the DNA by addition of 0.5 µl NT buffer per well. 1 µl well-specific indexing primer mix from Nextera XT Index Kit v2 Sets A-D and 1.5 µl NPM was added to each well and the tagmented cDNA was amplified for 14 cycles according to manufacturer’s specifications. PCR products from all wells were pooled and purified with 1× Agencourt AMPure XP beads (Beckman Coulter) according to the manufacturer’s protocol. The fragment size distribution was determined using a High Sensitivity DNA5000 assay for the Tapestation 4200 (Agilent) and library concentration was determined using a Qubit dsDNA HS assay (Thermo Fischer). Libraries were clustered at 1.4 pM concentration using High Output v2 chemistry and sequenced on a NextSeq500 system SR 75 bp with 2*8 bp index reads. Single-cell data were demultiplexed using bcl2fastq2 v2.20. and pseudo aligned to the mm10 mouse genome (Release M16 Gencode; GRCm38.p5) transcriptome using kallisto v0.44.0 (Bray et al., 2016). The sequences of all oligos used for SmartSeq2 can be found in the table submitted together with the raw data (GSE244474).

### scRNAseq analysis of CNS CD45^+^ cells

Unique Molecular Identifier (UMI) corrected expression matrices were imported into R as Seurat objects using Seurat v.4.0.2 (Hafemeister and Satija, 2019). From 407,803 barcodes and 28,441 genes, only protein coding genes were kept, reducing the number of genes to 17,942. Following the recommendations of Luecken and Theis (Luecken and Theis, 2019), 5,378 good quality cells were identified with 13,123 sufficiently expressed genes. After correcting for ambient genes using the SoupX automatic estimation by Young and Behjati (Young and Behjati, 2020), further reduction to 12,989 genes was achieved. 2,395 (11,935 genes) of the 5,378 cells were identified as *Spg15^-/-^* and 2,913 (11,512 genes) as *Ctrl*. After removing doublets using DoubletFinder (McGinnis et al., 2019), the total number of cells were reduced to 5,233.

Sequencing data were normalized, transformed and scaled using scTransform (Hafemeister and Satija, 2019). After performing Principal Component Analysis (PCA), the first 21 Principal Components (PCs) were used for Uniform Manifold Approximation Projection (UMAP) and construction of the Shared Nearest Neighbor (SNN) graph. The SNN graph was then used to perform clustering with the Louvain algorithm (Traag et al., 2019). For the clustering resolution, the results from Clustree v.0.4.3 (Zappia and Oshlack, 2018) and NbClust v.3.0 (Charrad et al., 2014) were taken into consideration. For these clusters the marker genes (one against all) were determined using the Wilcoxon rank sum test (Wilcoxon test) (Haynes, 2013a) with a 0.25 Log_2_-Fold-Change (Log_2_FC) cutoff and a threshold for a minimal of 10 % of cells expressing a marker gene in question. Canonical marker genes were used for cluster annotation. The protein expression matrix from the spiked-in TotalSeq antibodies was added to the processed Seurat object, in order to visualize differential protein expression between genotypes within the defined clusters.

### Gene Set Enrichment Analysis (GSEA)

Gene signatures for microglia were obtained from Hammond et al. (Hammond et al., 2019). A module score enrichment (Tirosh et al., 2016) was performed on our dataset using these signatures, filtering genes which are present in our expression matrix. 10 expression bins were used. In each bin 50 control genes were randomly selected as background to perform the signature enrichment against them. All cells with a module score of <=0 were removed to compare only cells with an enriched DAM signature. Then, the resulting module score was Log_2_ transformed and Log_2_FC was calculated for each *Spg15^-/-^* cell, comparing each module score to the median module score over all *Ctrl* cells.

For analysis of the functional phenotype in T cells of *Spg15^-/-^* mice the genes present in the dataset were ranked by their average expression in the respective genotype. This ranked gene list was then transformed from gene names into EntrezID using ClusterProfiler v.4.0.5 (Yu et al., 2012; Haynes, 2013b) and the Mouse annotation database org.Mm.eg.db v.3.12.0 (Carlson, 2020). Then, GSEA was performed using ClusterProfiler v.4.0.5 (Yu et al., 2012) to determine, if GO terms from the Gene Ontology (GO) (Biological Processes) database (Ashburner et al., 2000) were enriched in T cells from *Ctrl* or *Spg15^-/-^* animals, respectively. To test statistical significance of the enriched terms, a hypergeometric test (Yu et al., 2015) in combination with the Bonferroni procedure for multiple testing correction was performed (Yu et al., 2015, 2012; Haynes, 2013b). A cut-off of <0.05 for the adjusted p values was used to select significantly enriched GO terms.

### Gene Ontology Enrichment Analysis (GOEA)

First, all genes present after QC were transformed from gene names into EntrezID using ClusterProfiler v.4.0.5 (Yu et al., 2012; Haynes, 2013b) and the mouse annotation database org.Mm.eg.db v.3.12.0 (Carlson, 2020). All genes found in the GO database were used as background to test against when performing the hypergeometric test (Yu et al., 2015) for significance testing of the enriched term, in combination with the Bonferroni procedure for multiple testing correction (Yu et al., 2015, 2012; Haynes, 2013b). The Differential Expressed Genes (DEGs) between genotypes were determined by the Wilcoxon test using a 0.25 log_2_FC cutoff and a threshold for a minimum of 10 % of the cells expressing the gene in question. The enrichment was performed for terms of the Gene Ontology (GO) (Biological Processes) using ClusterProfiler v.4.0.5 (Ashburner et al., 2000; Yu et al., 2012). A cut-off of <0.05 for the adjusted p values was used to select significantly enriched GO terms.

### scRNAseq analysis of CNS T cells

The transcript abundance files in hdf5 format (Hoefling and Annau, 2020) from the pseudo alignment via kallisto v0.44.0 (Bray et al., 2016) were merged and converted into Transcripts per Kilobase Million (TPM) normalized gene expression matrices, with additional length scaling (”lengthScaledTPM”) and otherwise default parameters, via the package tximport v.1.18.0 (Soneson et al., 2015). The Ensembl IDs were transformed into gene symbols using biomaRt v.2.46.0 (Durinck et al., 2005, 2009). The expression matrix and respective metadata were added to a Seurat object. Additionally, fluorescence intensity values from the index sorting to 384 well plates were exported from the index sort fcs files as csv files according to the manufacturer’s recommendation (Penter et al., 2018) were added to the merged Seurat object with dplyr v.1.0.7 and rlist v.0.4.6.1 (Ren, 2016). Cells in the Seurat object were labeled as tdTomato^+^ or tdTomato^-^ based on the fluorescence intensity of this lineage tracer. The Seurat object was further preprocessed, by keeping only protein coding genes, and genes for the immune receptors (TCR and B Cell Receptor (BCR)). Ribosomal and genes which were predicted by genome assembly, but in most cases not associated with a function (gene names starting with “Gm”) were also removed. The above process resulted in 15,134 genes left from the initial 26,970 genes. Cells with less than 700 genes were removed from the analysis, resulting in 871 cells from the initial 1,471 cells. 526 cells remained for further analysis after removing CD3 negative cells. Gene expression was normalized to the library size per well, multiplied by 10^4^ and Log1p transformed, followed by centering and scaling (Hao et al., 2021). The PCA was calculated without truncated singular value decomposition (Hao et al., 2021; Li et al., 2019). The first 20 PCs were used for SNN and UMAP computation. The clustering was performed using the Louvain algorithm (Traag et al., 2019). The graph from the package Clustree v.0.4.3 (Zappia and Oshlack, 2018) and the different metrics from the package NbClust v.3.0 (Charrad et al., 2014) suggested a number of 9 clusters to be the most stable. Clusters were annotated using canonical markers. Differences in number of UMIs and contamination with *Olfr* genes hindered the annotation of T cell subsets, therefore a regression for these parameters was performed. For the regression of *Olfr* genes, a module score enrichment (Tirosh et al., 2016) was performed. Afterwards, ten clusters were found and annotated using canonical markers. The annotation correlated with protein expression levels from the index sort (Penter et al., 2018). After non-T cell clusters were removed, five T cell subsets were identified using analog and associated genes to the FACS panel for CNS T cells described above.

### TCR analysis

TCR sequences were extracted from fastq files generated with SmartSeq2 as described above using TRUST4 v.1.0.7 (Song et al., 2021). The resulting AIRR table (Vander Heiden et al., 2018) was combined with the transcriptome/protein expression matrix from above and analyzed using Scirpy v.0.11.1. Dominant clones were identified in *Spg15^-/-^* and *Ctrl* and a DEG test was performed. DEGs were used to perform GOEA as described above.

### RNA velocity of CNS T cells

Prior to calculating the velocity, unsorted Binary Alignment Map (BAM) (Li et al., 2009) files were generated from raw fastq files of each well from the SmartSeq2 protocol with STAR v.2.7.10b (Dobin et al., 2013). Splicing information was extracted from the unsorted BAM files using the docker image asaglam/biotools:21.01 in a Singularity runtime. Snakemake v.5.3.1 was applied to sort and index unsorted BAM files using Samtools v.1.9 (Li et al., 2009) and determine the splicing state of genes using Velocyto v.0.17.16 (La Manno et al., 2018). The resulting loom files contained two expression matrices for spliced and unspliced gene counts (La Manno et al., 2018; Bergen et al., 2020). These files were converted to Anndata objects, concatenated into one object and merged with the Anndata object containing the previously processed Seurat object, which was converted to Anndata using SeuratDisk v.0.0.0.9019. After filtering out CD4^+^ T cells and contaminating cells, 5000 top expressed genes and 20 minimally shared counts were selected for the calculation of velocity vectors. The neighborhood graph in Scanpy (Wolf et al., 2018) generated using the first 30 Principal Components (PCs) and 30 neighbors was taken for the calculation for the first and second order moments, thus accounting for the probabilistic nature of biological systems (Bergen et al., 2020; La Manno et al., 2018; Fröhlich et al., 2016).

A likelihood based dynamical model, considering individual splicing kinetics, was applied. Initial and terminal states and gene trends were inferred using CellRank v.1.5.0 (Lange et al., 2022).

### Trajectory analysis of microglial subtypes

Monocle3 v1.0.0 was used to infer developmental trajectory in the scRNAseq data of microglial subtypes. Detailed procedures were described by Cao et al. (Cao et al., 2019) the first 22 dimensions were used to generate a UMAP on microglia only. A connection matrix was then calculated using a modified partitioned approximate graph abstraction (PAGA) (Wolf et al., 2017) algorithm followed by significance testing of connections between clusters identified via PAGA and Louvain clustering. A principal graph on the low dimensional space was then learned using a modified SimplePPT algorithm (Qi Mao et al., 2017) Pseudotime was inferred using the node in the homeostatic microglia cluster, which had the highest distance in the UMAP to the intermediate microglia cluster, as the root node.

### Intercellular communication

The analysis of intercellular communication was performed according to the recommendations by Jin et al. (Jin et al., 2021) using CellChat v.1.1.3 (Jin et al., 2021) for the integrated dataset of microglia and T cell subsets. First, Seurat objects were subsetted for microglia subsets and T cells and integrated using Seurat (Butler et al., 2018). The integrated Seurat object was then split by the genotypes into *Ctrl and Spg15^-/-^*. All three Seurat objects (*Ctrl*, *Spg15^-/-^*, *Ctrl* & *Spg15^-/-^*) were converted into Cellchat objects using the normalized and log1p transformed expression matrices (Jin et al., 2021). The expression data was subsetted to keep only known signaling genes. Subsequently, overexpressed ligand and receptor genes were identified by the Wilcoxon rank sum test with a significance level of 0.05. The DEGs were then corrected for noise by calculating the average expression of a gene for each cell group, meaning that the quantiles of the gene expression were summed in a weighted manner. Jin et al. (Jin et al., 2021) based the prediction of gene-gene interaction on the Protein Protein Interaction (PPI) networks from STRINGDb (Szklarczyk et al., 2019), assuming physical interaction between the ligand and receptors, so that law of mass can be applied. For that purpose, they projected the expression profiles of the signaling genes onto the PPI by using the random walk network propagation (Cowen et al., 2017). Based on the weights derived from the networks, an interaction probability (e.g. strength) could be modeled. Statistically significant communication pathways between cell groups were identified by using the permutation test. Furthermore, the social network analysis tool sna (Butts, 2008) was applied to calculate the information flow between cell clusters using the metrics out-degree in-degree flow betweenness and information centrality.

### Quantification and statistical analysis

All statistical analyses, except for the analyses of sequencing data, were performed with GraphPad Prism software v5-8 (GraphPad Software). When analyzing statistical differences between two groups, two-tailed unpaired Student’s t-tests were performed. Statistical differences between three or more groups treated under similar conditions were analyzed by one-way ANOVA with Dunnett’s or Tukey’s multiple comparison tests. Two-way ANOVA were performed when comparing multiple groups in the context of different conditions. Sidak’s or Tukey’s multiple comparison tests were performed dependent on the experimental conditions. If the same cells or mice were measured at different time points, repeated measure analysis was performed. P values of less than 0.05 were considered to be significant (ns indicates not significant p > 0.05, p < 0.05 = *; p < 0.01 = **; p < 0.001 = ***). Data are representative of at least two independent experiments with at least 3 animals per group. Descriptive statistics, the performed statistical tests as well as the number of samples are stated in the figure legends.

### Code and data availability

Cleaning, dimensionality reduction, clustering, DEG testing and GOEA were performed on the docker image alefrol94/scrnaseq.analysis:reticulate. Any additional packages installed were tracked with the renv package v.0.14.0. Quantification and visualization of cell numbers/proportions and TCR analysis was done on the docker image wollmilchsau/Scanpy_scCODA:latest (from 4 November 2021). Velocity analysis was performed using the docker image nvcr.io/nvidia/pytorch:21.08-py3. Trajectory analysis of microglial subsets with Monocle3 (Cao et al., 2019) was performed using the docker image jsschrepping/r_docker:jss_R403_S4cran. Conda environments used for the analysis are saved as yaml files. The complete code can be found in the repository https://gitlab.dzne.de/frolova/spg15 and can be made available upon request. RNAseq and TotalSeq data are available at GEO under GSE244539.

---

**Figure S1.**
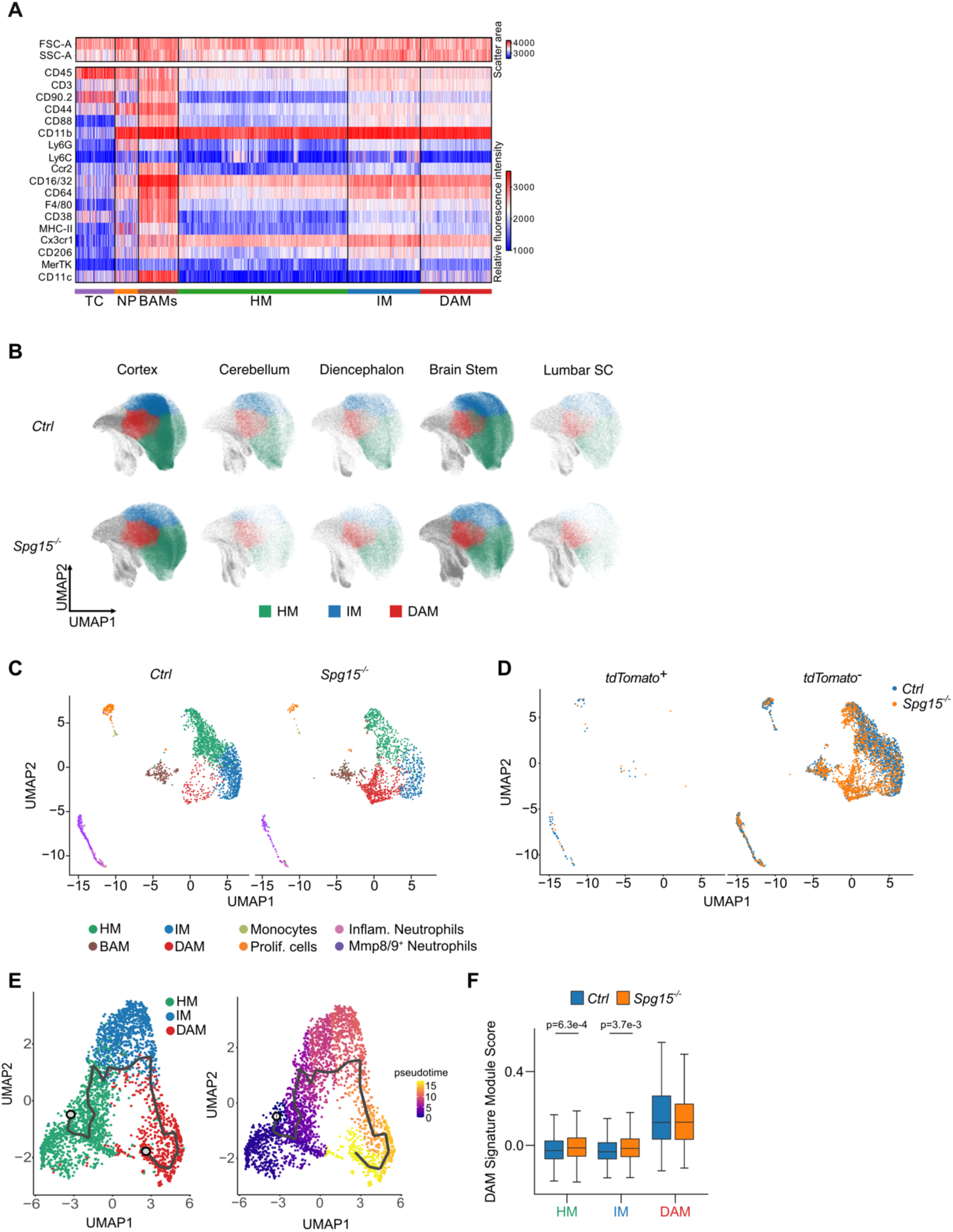
Immunophenotyping of microglia in the CNS of *Spg15^-/-^* mice. **A.** Fluorescence intensity in a linear (SSC/FSC) or biexponential scale (rest) of surface marker expression on single cells shown in Fig. 2A. **B.** UMAPs showing microglia clusters (homeostatic, HM; intermediate, IM; disease-associated, DAM) for single CNS regions from *Spg15^+/+^; Cxcr4^CreERT2^; Rosa26^tdTomato^ (Ctrl)* and *Spg15^-/-^; Cxcr4^CreERT2^; Rosa26^tdTomato^* (*Spg15^-/-^*) mice. Combined clusters are depicted in Fig. 2A. **C.** UMAPs of scRNA-seq data from old *Ctrl* and *Spg15^-/-^* brains shown combined in Fig. 3A. **D.** UMAP from Fig. 3A split into tdTomato^+^ and tdTomato^-^ cells. **E.** Trajectory analysis of microglia using Monocle3. The trajectory path from HM to DAM is superimposed on the UMAP generated on microglia only (left). UMAP colored by pseudotime shows the distance to the defined starting node (labeled as 1) in the HM cluster (right). **F.** The module score for a DAM gene signature was calculated for each cell in the microglia subclusters. The distribution of module scores for each microglia subcluster was visualized using a boxplot. Outer boundaries represent the 75- or 25-percentile, respectively. Middle line represents the median. Whiskers indicate extreme values 1.5 times the interquantile range smaller or larger than the respective percentile. A double sided wilcoxon rank sum test in combination with the Holm method for multiple test correction was used to test differential enrichment of the DAM signature in *SPG15^-/-^* vs. *Ctrl* animals for each microglia subcluster.

**Figure S2.**
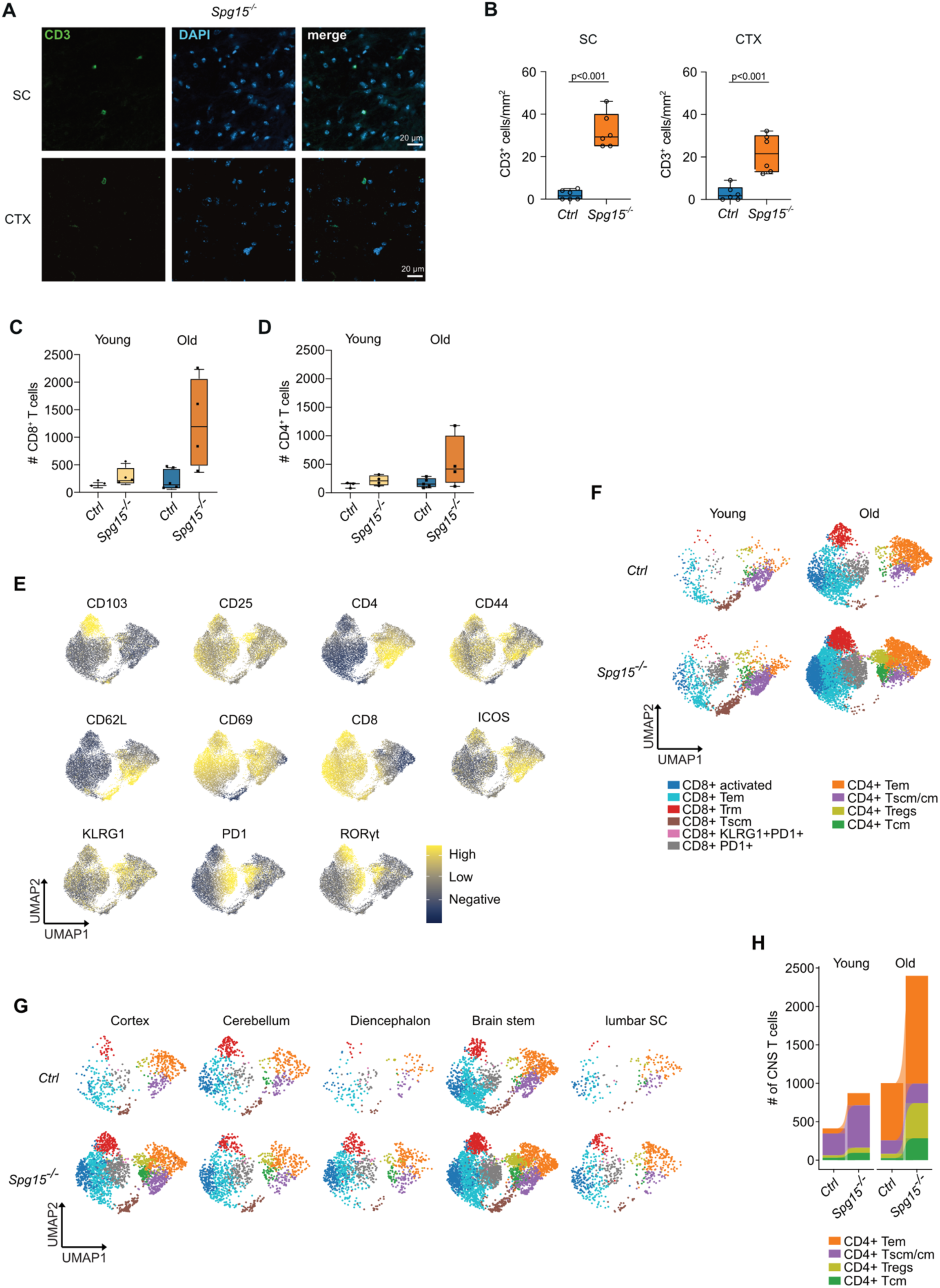
Analysis of the enriched T cell compartment in the CNS of *Spg15^-/^*^-^ mice. **A.** Representative IF staining for CD3 in spinal cord (SC) and cortex (CTX) from 12-15 month-old *Spg15^-/-^; Cxcr4^CreERT2^; Rosa26^tdTomato^* (*Spg15^-/-^*) mice. **B**. Quantification of CD3^+^ cells per mm^2^ in SC and CTX of *Ctrl* and *Spg15^-/-^* mice aged 12 – 15 months. **C, D.** Number of CD8^+^ (C) and CD4^+^ (D) T cells in the CNS of young (2-3 months) and old (12-15 months) *Ctrl* and *Spg15^-/-^* mice analyzed by flow cytometry. **E.** Protein marker expression used for UMAP generation and cluster annotation in Figure 4E. **F.** Distribution of T cells after splitting the UMAP from Figure 4E according to genotype and age. **G.** UMAPs showing T cells from old animals after splitting the UMAP from Figure 4E according to genotype and brain region. **H.** Compositional changes of CD4^+^ T cell subtypes as defined in Figure 4E. **B, C, D** Statistical significance was assessed with an unpaired Student’s *t*-test.

**Figure S3.**
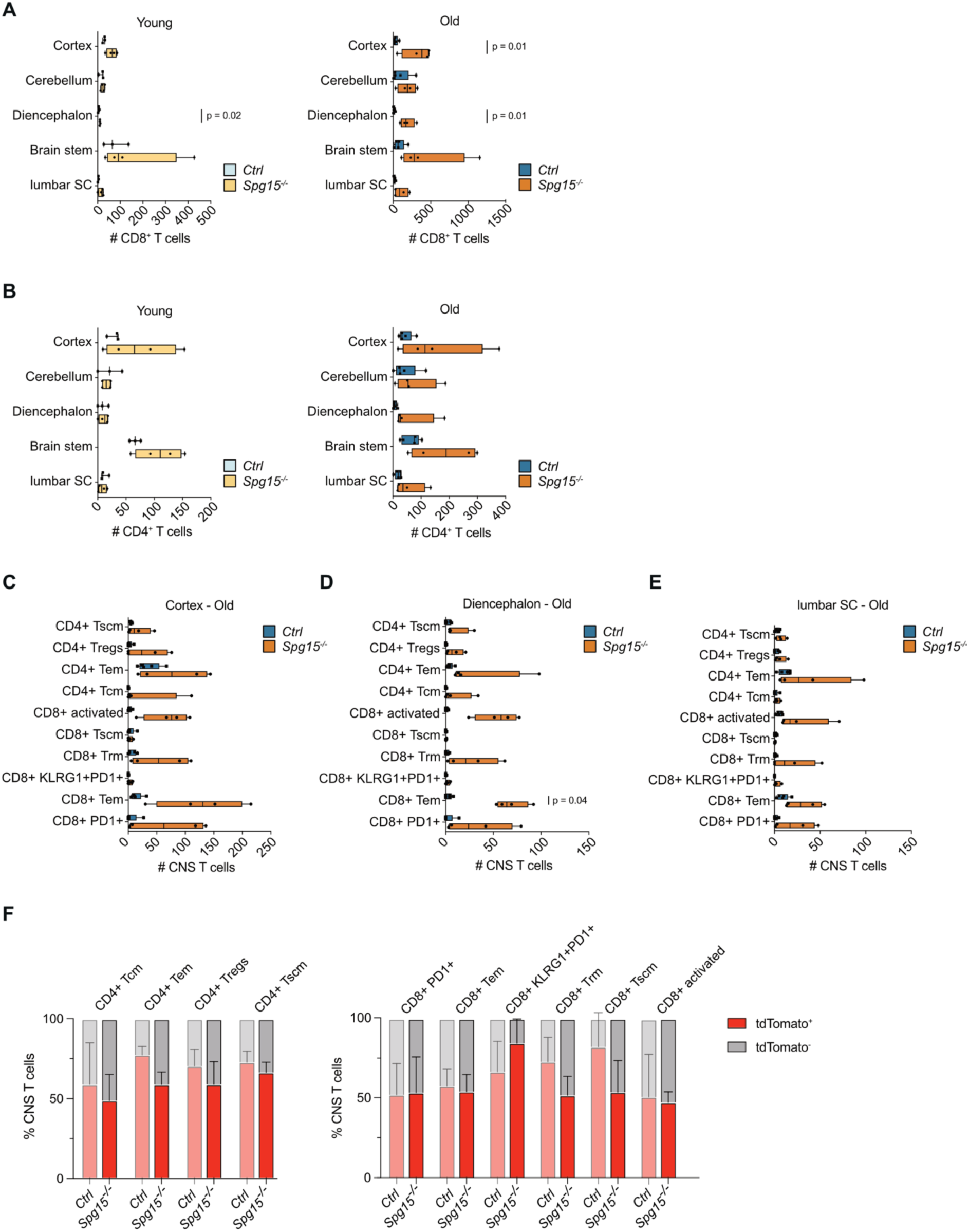
High-dimensional flow cytometry analysis of the T cell compartment in the CNS of *Spg15^-/^*^-^ mice. **A,B.** Number of CD8^+^ (**A**) and CD4^+^ (**B**) T cells in various brain regions of young and old *Spg15^+/+^; Cxcr4^CreERT2^; Rosa26^tdTomato^ (Ctrl)* and *Spg15^-/-^; Cxcr4^CreERT2^; Rosa26^tdTomato^* (*Spg15^-/-^*) mice. **C, D, E.** Number of T cells per T cell subset in cortex (**C**), diencephalon (**D**), and lumbar SC (**E**). For definition of T cell subsets see Figure 4E. **F.** tdTomato expression in CD3^+^ T cell subclusters present in the CNS of old (12-15 months) *Ctrl* and *Spg15^-/-^* mice. **Left:** Subclusters found in CD4^+^ T cells. **Right:** Subclusters found in CD8^+^ T cells. Data is represented as mean with SEM. **A** Statistical significance was assessed with an unpaired Student’s *t*-test. **C, D, E** Statistical significance was assessed with an one-way FDR-corrected ANOVA.

**Figure S4.**
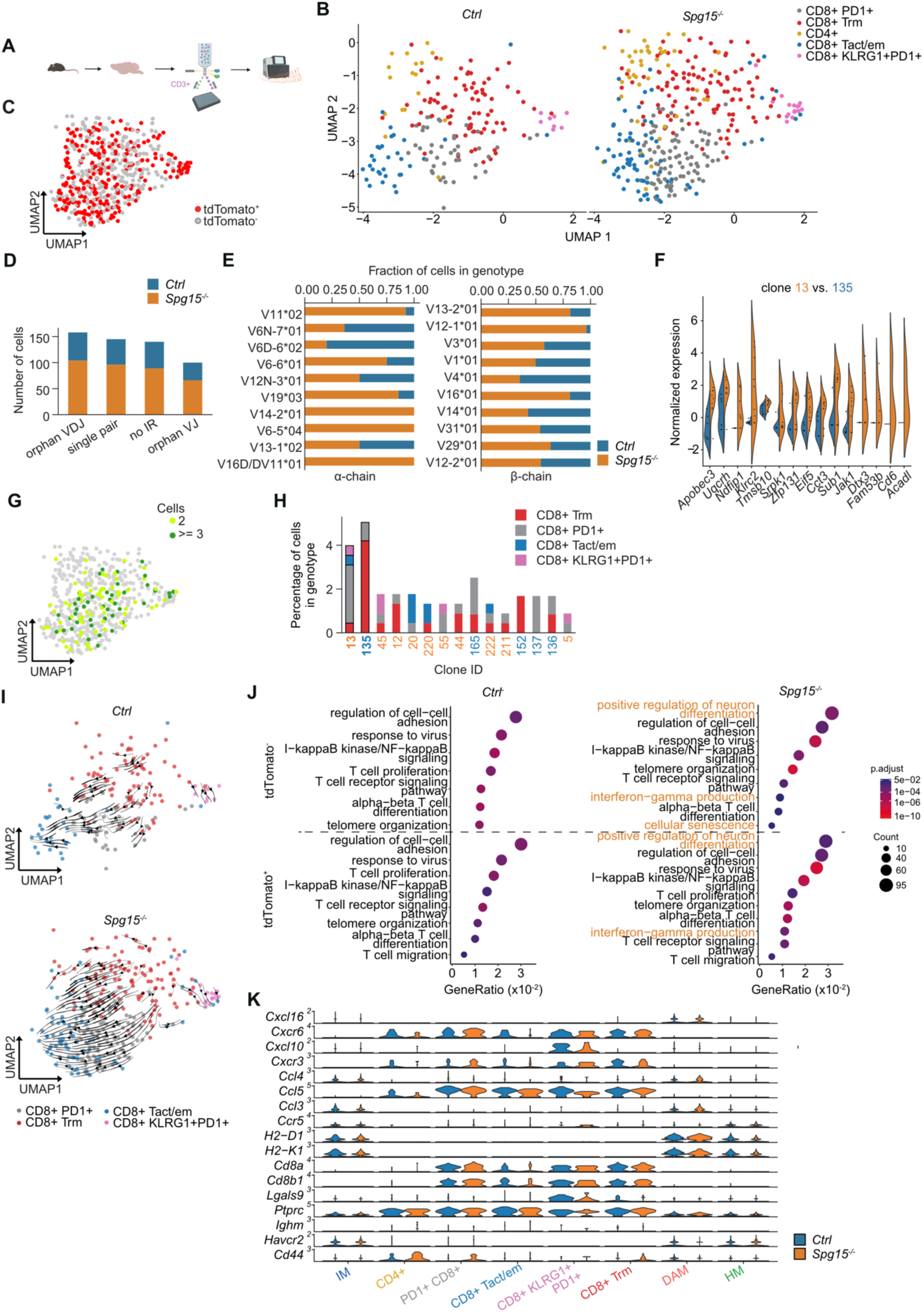
TCR analysis and tdTomato tracing confirms clonally independent expansion of resident CNS CD8^+^ T cells in the CNS of old *Spg15^-/-^*mice. A. Workflow for the generation of full length scRNA-seq libraries with the SmartSeq2 chemistry from CNS T cells isolated from old (12-15 months) *Spg15^+/+^; Cxcr4^CreERT2^; Rosa26^tdTomato^ (Ctrl)* and *Spg15^-/-^; Cxcr4^CreERT2^; Rosa26^tdTomato^* (*Spg15^-/-^*) animals. Created with Biorender. **B.** UMAP from full length scRNA-seq data from CNS T cells isolated from old *Ctrl* and *Spg15^-/-^*animals split by genotype (n = 2 per genotype). **C.** tdTomato expression in single T cells was mapped onto the UMAP from Figure 5. **D.** Quality control of the TCR sequences recovered from the SmartSeq2 full length libraries using TRUST4 split into *Ctrl* (blue) and *Spg15^-/-^* (orange). Cells were grouped according to the following information: orphan VDJ: TCR β-chain only; single pair: TCR α-chain and β-chain; no IR: No TCR inferred; orphan VJ: TCR α-chain only. **E.** Percentage of clones from *Ctrl* (blue) or *Spg15^-/-^* animals (orange) using the respective V gene segment of the TCR α-chain gene locus (left) or β-chain gene locus (right). **F.** Expression levels of the top 15 DEGs between the dominant clone in *Ctrl* (135) and *Spg15^-/-^* animals (13). **G.** Expanded clones were mapped onto the UMAP from Figure 5 to visualize transcriptional similarity of expanded clones. **H.** Compositional analysis of clones. Percentage of the clones from the respective genotype was visualized on the x-axis. The cell cluster identity was color coded. Dominant clones in *Ctrl* (Clone ID 135, blue) and *Spg15^-/-^* animals (Clone ID 13, orange) were highlighted. **I.** Velocity vectors were calculated for T cells from control (left) and *Spg15^-/-^*^-^ animals (right) and superimposed on the split UMAP from Figure 5. **J.** Enriched GO terms by GO enrichment analysis using DEGs between either tdTomato^+^ (top) or tdTomato^-^ CD8^+^ T cells (bottom) from *Ctrl* and *Spg15^-/-^* animals. Terms uniquely enriched in *Spg15^-/-^* animals were highlighted in orange. **K.** Normalized expression of inferred ligand receptor pairs by CellChat visualized as violin plots for microglia and T cell subsets. Blue color indicates expression in *Ctrl* and orange color in *Spg15^-/-^* animals.

